# Synergistic induction of a lethal Autosis-to-Apoptosis switch by Phytocannabinoids and beta-Caryophyllene in Triple-Negative Breast Cancer Cells

**DOI:** 10.64898/2026.04.05.716056

**Authors:** Carla Hamann, Olivia Jansen, Kateline Jullien, Liza Lhonneux, Allison Ledoux, Michel Frédérich, Erik Maquoi

## Abstract

**Background:** Triple-negative breast cancer (TNBC) presents significant therapeutic limitations due to its aggressive heterogeneity and the rapid emergence of adaptive resistance to apoptosis-based regimens. Addressing these challenges requires polypharmacological strategies capable of modulating multiple signalling networks simultaneously. While the *Cannabis sativa* phytocomplex offers a vast chemical space for multi-target intervention, the quantitative pharmacological basis of its synergistic interactions remains largely uncharacterised.

**Purpose:** This study aimed to deconstruct the synergistic landscape of high-purity phytocannabinoids (CBD, CBG, CBD-A) in combination with the sesquiterpene β-caryophyllene (BCP) against TNBC, using MDA-MB-231 as a primary model and Hs578T as a validation line.

**Methods:** Growth Rate (GR) inhibition metrics and the SynergyFinder+ framework were used to map pharmacological interactions across four reference models. Subcellular dynamics and phenotypic transitions were characterised by high-resolution label-free holotomographic microscopy combined with live-cell kinetic imaging and single-cell fate mapping.

**Results:** Two highly potent synergistic clusters were identified for CBD-CBG-BCP combinations, with ZIP, HSA, and Bliss synergy scores exceeding 65. CBD-A exhibited minimal interaction potential and was excluded from ternary studies. GR-based quantification further revealed that these combinations produced net cytotoxicity (GR < 0) at sub-IC₅₀ concentrations of each component. Single-cell fate mapping by holotomographic microscopy identified a temporally ordered death programme: an initial phase of extensive cytoplasmic vacuolisation associated with focal perinuclear space swelling and progressive nuclear compression, morphological hallmarks of autosis, which is followed by a transition to apoptotic execution. The autotic nature of the primary death phase was confirmed by pharmacological rescue with digoxin, a selective inhibitor of the Na⁺,K⁺-ATPase. To the best of our knowledge, this sequential engagement of autosis followed by apoptotic execution represents the first documented instance of such a two-stage death programme in any cellular model.

**Conclusion:** These findings provide robust evidence that specific phytocannabinoid-terpene ratios engage a Na⁺,K⁺-ATPase-regulated autotic programme as an upstream commitment step, followed by apoptotic execution, effectively circumventing the caspase-independent resistance mechanisms characteristic of TNBC. This study establishes a rational, quantitatively validated framework for transitioning from empirical botanical use to evidence-based, multi-target cannabinoid polypharmacology in aggressive breast cancer.

## 1. Introduction

Plant extracts have long been recognised as a rich source of bioactive compounds with significant therapeutic potential. These complex mixtures contain diverse phytochemicals that may interact through multiple mechanisms to produce biological effects greater than those predicted by individual constituents [1]. While contemporary drug discovery has historically favoured “single-target, single-compound” models, ethnopharmacological evidence and modern network pharmacology increasingly support the use of multi-component preparations to address the adaptive resistance mechanisms of complex malignancies [2–3]. The combination of compounds represents a strategic advantage in oncology, enabling dose reduction, minimising off-target toxicities, and overcoming tumour heterogeneity by targeting multiple sites of action [4–5]. To evaluate the potential of combinations, drug-drug interactions are rigorously classified into synergy, additivity, or antagonism, requiring robust computational models to validate the therapeutic gain of any combined formulation [6]. Synergy occurs when the combined effect of multiple compounds exceeds the sum of their individual effects, additivity arises when the combined effect equals the expected sum of individual effects, and antagonism is observed when the combined effect is less than expected [7].

A compelling model for studying such polypharmacological interactions is *Cannabis sativa* L., which possesses a rich phytochemical profile comprising more than 120 cannabinoids, alongside a diverse array of terpenes and flavonoids [8–9]. While Δ^9^-tetrahydrocannabinol (THC) and cannabidiol (CBD) have received considerable attention for their therapeutic properties, there is growing consensus that the medicinal potential of Cannabis arises from its phytocomplex. Historically described as the “entourage effect” [10], this phenomenon is now understood as a polypharmacological synergy, where minor cannabinoids and terpenoids act as pharmacological partners that modulate the pharmacodynamics of primary constituents.

Despite the widespread interest in these interactions, rigorous scientific evidence quantifying synergy remains limited, with many studies failing to distinguish between true synergy and simple additivity. To advance the therapeutic application of non-psychotropic cannabinoids, this study aims to evaluate potential synergistic interactions between several prevalent compounds: CBD, cannabigerol (CBG), cannabidiolic acid (CBD-A) and β-caryophyllene (BCP), the most abundant sesquiterpene in hemp. This evaluation focuses specifically on their combined efficacy against triple-negative breast cancer (TNBC) cells, a subtype for which effective therapeutic options remain urgently needed. TNBC is characterised by high inter-tumoral heterogeneity and the absence of oestrogen, progesterone, and HER2 receptors, limiting the success of conventional targeted therapies and leading to poor clinical outcomes. Recent experimental evidence has demonstrated that cannabinoids can bypass these limitations by modulating critical pathways in TNBC, including the induction of autophagy, ER-stress-mediated apoptosis, and the inhibition of the AKT/mTOR axis [11–13].

Among the selected compounds, BCP stands out as a selective agonist of the cannabinoid receptor CB2 [14] and has garnered significant interest due to its demonstrated therapeutic potential, including anti-inflammatory, analgesic, and anticancer properties [15–16].

By utilizing advanced high-content imaging and robust interaction models (Bliss, Loewe, HSA, ZIP), this research provides a systematic understanding of phytocannabinoid-terpene synergies. These findings are essential for guiding the rational development of optimised cannabis-based therapeutic formulations.

## 2. Materials and methods

### 2.1. Chemicals

Cannabidiol (#85705), cannabigerol (#85956), and cannabidiolic acid (#85839) were purchased from PhytoLab (Germany) with a purity ≥ 99.0 % (HPLC). β-caryophyllene (≥ 98.0% GC; #22075), doxorubicin, digoxin (#D6003) and staurosporine (#S6942) were purchased from Sigma-Aldrich. Stock solutions (5 mg/mL for cannabinoids and 12 mg/mL for BCP) were prepared in dimethyl sulfoxide (DMSO). Cannabinoid solutions were stored at-20°C, while BCP solutions were freshly prepared for every test. The final DMSO concentration was maintained at a maximum of 0.5% across all conditions.

### 2.2. Cell culture

Human TNBC cell line MDA-MB-231 and Hs578T was kindly provided by the Laboratory of Tumour and Development Biology (GIGA institute, Liège, Belgium) and maintained in high-glucose Dulbecco’s Modified Eagle Medium (DMEM) supplemented with 10% fetal bovine serum (FBS), 100 IU/mL penicillin, 100 µg/mL streptomycin, 1 mM sodium pyruvate, and 2 mM L-glutamine. All culture reagents were purchased from Thermo Fisher Scientific. Cells were maintained in a humidified atmosphere at 37°C with 5% CO_2_.

### 2.3. Cell vitality assessment using metabolic-based assay

MDA-MB-231 (2500 cells/well) and Hs578T (1500 cells/well) lines were seeded in 96-well plates (100 µL) and allowed to adhere for 24 h prior to treatment. Test compounds or combinations (50 µL) diluted in culture medium were added in triplicate to the plate. Doxorubicin (5 µM) was used as a reference standard for positive efficacy control in all assays. For experiments containing BCP, plates were sealed with a breathable membrane (#Z380059, Merck) to prevent evaporation and cross-contamination between surrounding wells. After 72 hours of incubation, the medium was removed and replaced with 75 µL of a 10 times diluted solution of PrestoBlue reagent (#A13262, Invitrogen^TM^, Thermo Fisher Scientific). After 2 hours of incubation at 37°C, fluorescence was measured using a FlexStation 3 microplate reader (Molecular Devices, USA) with excitation and emission wavelengths of 560 nm and 590 nm, respectively.

### 2.4. Live-cell imaging and image quantification

Live-cell imaging was performed to analyse the cellular phenotypic alterations induced by hemp-derived compounds on cell populations over time. Two different imaging modalities were used: phase contrast microscopy associated with fluorescence imaging using an Incucyte® SX5 Live Cell Imaging System (Sartorius, Germany) and a label-free holotomographic microscopy (HTM) using a CX-A system (Nanolive, Switzerland).

For the Incucyte® SX5 system, imaging was performed with a 10x objective in phase contrast, as well as in green (Ex: 453-485nm; Em: 494-533nm), orange (Ex: 546-568nm; Em: 576-639 nm) and near-infrared (NIR - Ex: 648-674nm; Em: 685-756nm) fluorescence channels. Images (4 fields per well) were acquired kinetically every 2 hours for a total period of 72 hours, or at initial and final timepoints (0 and 72h, respectively), depending on the experiment design. Cell nuclei were labelled with SPY650-DNA (diluted 1/4000; Spirochrome, Switzerland), while dead cells were labelled using SYTOX Green (20 nM; Thermo Fisher Scientific) or SYTOX Orange (10 nM; Thermo Fisher Scientific), depending on the multiplexing requirements. To evaluate apoptosis, the NucView® 488 Caspase-3 enzyme substrate (0.5 µM; Biotium, USA) was used, with staurosporine (0.5 µM) as a specific positive control for caspase activation. For digoxin experiments, cells were pre-incubated with digoxin (0.5 or 1 µM) for 2 hours prior to the addition of treatments.

As BCP is volatile, 96-well plates were sealed with imaging compatible membranes (ibiSeal #10874, Ibidi, Germany). Automated image analysis routines were optimised for each label using the Incucyte SX5 software package (version 2024B) and training data from vehicle and compound-treated samples. Images were analysed using a routine with the following settings to count SPY650-DNA^+^ objects (Segmentation: Fixed threshold; threshold adjustment: 0.2; Edge split on; Edge sensitivity 0), SYTOX Green^+^ objects (Segmentation: Surface Fit; threshold adjustment: 1.5), SYTOX Orange^+^ objects (Segmentation: Top-Hat; radius 50; threshold: 1.0; Edge split on; Edge sensitivity 25), and Nucview^+^ object (Segmentation: Top-Hat; radius 50; threshold: 0.8; Edge split on; Edge sensitivity 0). To account for differences in proliferation rates between conditions, drug responses were also quantified using the Growth Rate (GR) metric, calculated by comparing the number of living cells (SPY650-DNA^+^ and SYTOX Green^-^ nuclei) in treated and control conditions at the same timepoints [17].

For HTM analysis, MDA-MB-231 cells (5×10^4^/mL) suspended in DMEM supplemented with 10 % FBS were seeded in a 50 µL droplet in the centre of 35-mm dish (Ibidi, #80136). Once adherent, 550 µL of medium was added and dishes were incubated overnight at 37°C in 5% CO_2_. Two-fold concentrated treatment solutions (600 µL, containing a maximum final DMSO concentration of 0.3%) were then added. Image acquisition started after a 2-hour equilibration period in the microscope chamber. Live-cell imaging was performed using CX-A® holotomographic microscope. This system is equipped with an incubator chamber that maintains samples under physiological conditions (37 °C, 5% CO_2_, 90% relative humidity). Image acquisition was conducted using a 60× dry objective, a 520 nm laser (irradiance: 0.2 nW/µm²; acquisition time: 45 ms per frame), and a CMOS camera (Blaser ace acA1920-155um). A total of 49 fields of view (7×7 grid scan) per 35-mm dish were captured every 20 minutes for 20 hours and processed using the Eve software (Nanolive).

### 2.5. Image analysis

The kinetic progression of individual cell phenotypes was analysed via lineage tracking of holotomographic datasets. Trajectories were generated and curated using the Mastodon plugin for Fiji [18], allowing for the discrete state-mapping of the autosis-to-apoptosis transition.

### 2.6. Drug combination assays

Cannabinoids (CBD, CBG and CBD-A) and BCP were first tested individually for their activity on the vitality of MDA-MB-231 cells and their IC_50_ values were determined using nonlinear regression analysis with a four-parameter logistic model (variable slope) in GraphPad Prism software (version 8.0). Each concentration was tested in triplicate with at least three independent experiments. To explore interactions among hemp-derived compounds and validate the entourage effect, various cannabinoid and cannabinoid–BCP combinations were assessed in MDA-MB-231 cells with cell vitality used as a readout. Data were expressed as the percentage of growth inhibition (drug vs control).

To assess the combinatorial effects of CBD with other cannabinoids or BCP, three different two-drug combinations were tested: (1) CBD and CBG, (2) CBD and CBD-A, and (3) CBD and BCP. Concentration ranges were selected based on initially determined IC_50_ values, including concentrations exceeding IC₅₀ to detect potential antagonistic effects (**Tables S1-3**).

A three-compound system (CBD, CBG, and BCP) was also designed to evaluate synergistic potential for mimicking the entourage effect in hemp. For this experiment, 4 concentrations each of CBD (0-16 µM), CBG (0-28 µM) and BCP (0-98 µM) were combined, resulting in a total of 64 unique combinations (**Table S4**). Vehicle-treated cells received culture medium supplemented with 0.3% DMSO.

Synergy analysis was performed using SynergyFinder+ software (http://www.synergyfinder.org) [19]. Synergy scores were calculated based on four reference models: Bliss, Loewe, HSA, and ZIP. The two-drug and three-drug combination functions of the software were used with no baseline correction mode.

### 2.7. Statistical analysis

All data were analysed using GraphPad Prism 8.0 (GraphPad software, USA). Results are presented as mean ± standard deviation (SD) or as median values, depending on the specific data distribution and visualization requirements of each figure. Each experiment was performed at least in triplicate across three independent experiments. Statistical comparisons were performed using one-way ANOVA (conducted on endpoint value or on area under the curve (AUC) for kinetic studies), followed by Tukey’s post hoc test for multiple comparisons. For drug combination studies, selected combinations were statistically compared to their corresponding individual drug treatments and vehicle control. For normalized data where the reference group was set to 100%, a one-sample t-test was performed to evaluate significant deviations from the baseline. Statistical significance was set at *p* < 0.05.

## 3. Results

### 3.1. Hemp-derived compounds differentially affect MDA-MB-231 cell vitality

We first examined the effects of individual cannabinoids and BCP on MDA-MB-231 cell vitality using a metabolic-based assay after 72 hours of treatment (**Fig. 1A**). CBD demonstrated the highest inhibitory activity (IC_50_ of 17.4 ± 1.7 µM), followed by CBG (IC_50_ of 28.1 ± 0.3 µM), representing the most bioactive cannabinoids in non-psychotropic cannabis varieties [20]. In contrast, minor cannabinoids such as CBD-A, the naturally occurring acidic precursor of CBD, exhibited a significantly lower activity (IC_50_ of 104.4 ± 12.6 µM). BCP showed the lowest activity (IC_50_ of 141.2 ± 24.6 µM). Given its established use as a first-line drug in TNBC chemotherapy, doxorubicin was used as a positive control, achieving over 95% inhibition of cell vitality at 5 µM in all independent tests.

**Figure 1:**
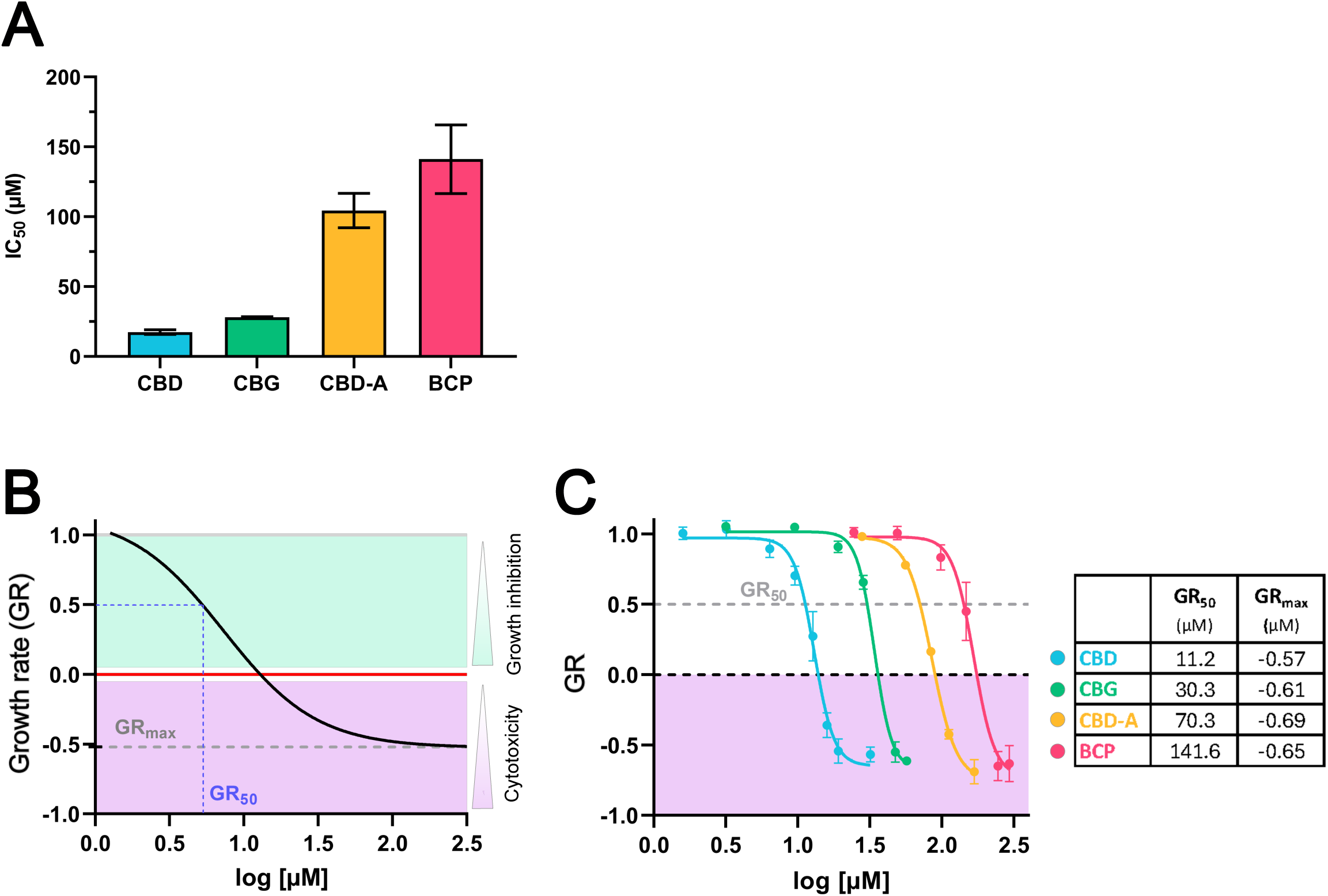
Hemp-derived compounds differentially affect MDA-MB-231 cell vitality and growth. MDA-MB-231 cells were treated with hemp-derived compounds: cannabidiol (CBD), cannabigerol (CBG), cannabidiolic acid (CBD-A), and β-caryophyllene (BCP). **(A)** IC_50_ values represent the concentration (µM) required to reduce cell vitality by 50% after 72 hours of treatment, as determined by resazurin-based metabolic assay. Data are expressed as the mean ± SD from four independent experiments performed in triplicate. Control conditions: 0.3% DMSO. **(B)** Illustration delineating the metrics of growth rate (GR), with GR>0 denoting a growth inhibitory effect, GR=0 indicating cytostasis, and GR<0 signifying cytotoxicity. GR_50_ and GR_max_ are projected onto the x-axis and y-axis, respectively. **(C)** GR dose–response curves at 72 hours of treatment for cells treated with the four compounds. GR values calculations were based on the number of SPY-DNA^+^/SYTOX Green^-^ nuclei quantified by live-cell imaging. GR_50_ and GR_max_ values (µM) for each compound are indicated in the table. Data represent mean ± SD from three independent experiments.

As metabolic-based assays indirectly reflect the number of living cells, a direct counting of the number of viable cells (SPY-DNA^+^/SYTOX Green^-^ nuclei) was performed by live-cell imaging after 0 and 72h of treatment. Results were expressed as drug-induced growth rate (GR) inhibition [17]. GR values range from +1 (no anti-proliferative effect) to-1 (complete cell death), with 0 reflecting a cytostatic activity, as illustrated in **Fig. 1B**. The GR_50_ represents the concentration reducing growth rate by 50%, reflecting the potency of the treatment.

As shown in **Fig. 1C**, CBD displayed the lowest GR_50_ value (11.2 µM), followed by CBG (30.3 µM), while CBD-A and BCP exhibited significantly weaker activities (70.3 µM and 141.6 µM, respectively). These results validate the IC_50_-derived potency ranking, establishing CBD and CBG as the most effective growth inhibitors. The GR_max_ (Growth Rate maximal effect) represents the maximum effect of the treatment at the highest tested concentration, reflecting its efficacy. Analysis of GR_max_ values demonstrated that the 4 compounds were highly cytotoxic when used at the highest concentrations tested. These differential activities serve as a prerequisite for subsequent synergy studies, which will assess if less active compounds, such as CBD-A, contribute to overall therapeutic efficacy via combinatorial effects.

### 3.2. Evaluation of synergistic interactions between phytocannabinoids and terpenes

Here, we utilized high-purity (>99%) phytocannabinoids and terpenes to deconstruct the “entourage effect” into quantifiable pharmacological interactions. Three binary combinations mixing neutral and acidic cannabinoids as well as cannabinoids and terpenes (1: CBD + CBG, 2: CBD + BCP and 3: CBD + CBD-A) were tested for their activity on MDA-MB-231 cell vitality using a metabolic-based assay. CBD + CBG combination achieved 50% inhibition at relatively low concentrations of each drug (8.0 and 19.0 µM, respectively), with high inhibitory effects across most concentrations (**Fig. 2A**). CBD + BCP showed similar trends from 6.4 μM and 97.9 μM (**Fig. 2B**). In contrast, CBD + CBD-A required substantially higher concentrations of both drugs to reach significant inhibition, with only the two highest doses of CBD-A (83.7 and 111.6 μM) showing strong activity (**Fig. 2C**).

**Figure 2:**
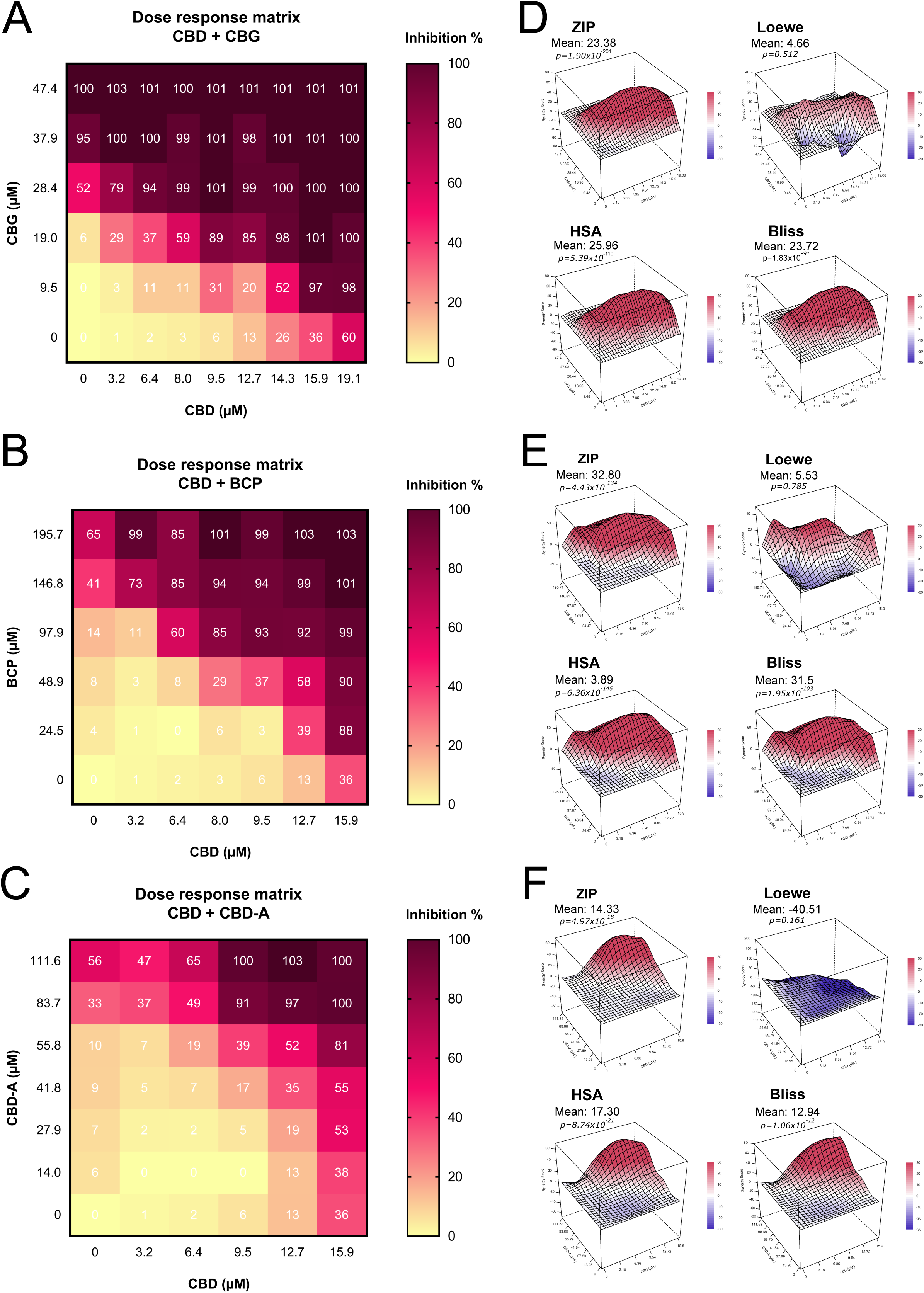
Analysis of drug interactions in pairwise combinations of hemp compounds on the vitality of MDA-MB-231 cells. (A-C) Dose-response heatmaps showing percentage inhibition in vitality assay after 72 hours of treatment with combinations CBD + CBG **(A)**, CBD + BCP **(B)** and CBD + CBD-A **(C)**. Colour intensity represents the percentage of inhibition relative to vehicle control (0.5% DMSO). **(D-F)** Three-dimensional synergy landscapes generated using SynergyFinder+ with ZIP, Loewe, HSA and Bliss reference models: CBD + CBG **(D)**, CBD + BCP **(E)**, CBD + CBD-A **(F).** Positive scores indicate synergy; negative values indicate antagonism. Data are presented as median from three independent experiments.

The potential synergy of compound combinations was analysed by using SynergyFinder^+^ [19]. Synergy scores were calculated using four established models to interpret interaction effects: Bliss, Loewe, highest single agent (HSA) and zero interaction potency (ZIP). Each model determines synergy, additivity, or antagonism by comparing experimentally observed effects with expected effects. The HSA model assumes that the expected effect equals that of the most active single agent at the given concentration. This model is particularly relevant when one compound exhibits minimal activity (e.g., CBD + CBD-A) [21]. The Bliss independence model assumes that drugs act independently on distinct, non-interfering targets but contribute to the same observed effect [7,21]. The Loewe model, based on dose equivalence principles, is most relevant for drugs with similar mechanisms and comparable dose-response curves and maximum effects. This makes it most relevant for combinations of CBD and CBG, as well as those involving CBD-A. The ZIP model was recently developed to overcome biases in existing models. It assumes that one drug’s addition does not alter the other’s potency or efficacy, regardless of whether they target the same or different pathways [22]. Synergy scores below –10 indicate antagonism, between –10 and +10 indicate additive effects, and above +10 indicate synergistic interactions [19]. Different models were used because these compounds could act on many potential targets, and no single model can fully capture all possible interactions or mechanisms of synergy.

Analysis of synergy scores (**Fig. 2D-F**) revealed that CBD + BCP was the most synergistic combination (mean scores: 32.8 ZIP, 31.5 Bliss), followed by CBD + CBG (23.4 ZIP, 23.7 Bliss) and CBD + CBD-A (14.3 ZIP, 12.9 Bliss). HSA model results were similar to ZIP and Bliss, reflecting the dominant contribution of CBD. In contrast, the Loewe model yielded lower synergy scores, suggesting additivity for CBD + BCP (5.5) and CBD + CBG (4.7), but indicating strong antagonism for CBD + CBD-A (–40.5). This model may overestimate antagonism when one compound exhibits a substantially lower contribution (**Fig. 2F**).

To complement the vitality assays, live-cell imaging of cancer cells was used to further investigate the potential antagonism between CBD and CBD-A (**Fig. 3**). While the combinations of CBD used at a sub-cytotoxic concentration (6.4 µM; < half of IC₅₀) with CBD-A (14.0–55.8 µM) resulted in negative Loewe synergy scores (–21.8 to –16.8, **Table S2**), no significant changes in GR values were observed compared with single compound treatments (GR=0.90 for CBD and 0.78 for CBD-A, **Fig. 3A-B**), indicating an apparent lack of synergy. These combinations exhibited limited cytotoxicity, as evidenced by percentage of the cell death which remains equivalent to that of the vehicle-treated cells, and not significatively different from the single treatments (**Fig. 3C-D**). This was corroborated by the presence of a similar number of SYTOX Green^+^ dead cells as in the control (compare **Fig. 3I** and **J**). The Bliss model provided modest synergy scores (–8.9 to +5.2, **Table S2**), consistent with the limited antiproliferative potency of CBD-A and CBD at these concentrations. When higher concentrations of CBD (12.7 µM; ≈ ¾ of IC₅₀) were combined with 41.8 or 55.8 µM of CBD-A, a transition from cytostatic to slightly cytotoxic effects were observed, with GR values of-0.01 and-0.29, respectively (**Fig. 3E-F**). This latter combination with 55.8 µM CBD-A (Bliss score: 30.9) significantly outperforms single compounds’ activities (**Fig. 3E-F, Table S2**). This observation was corroborated by a significant reduction in cell density associated with an increased cell death (compare **Fig. 3I** and **K**). These results indicate that only the highest dose of CBD-A enhances CBD efficacy. Consequently, due to its low potency-to-synergy ratio, CBD-A was excluded from subsequent ternary combination assays.

**Figure 3:**
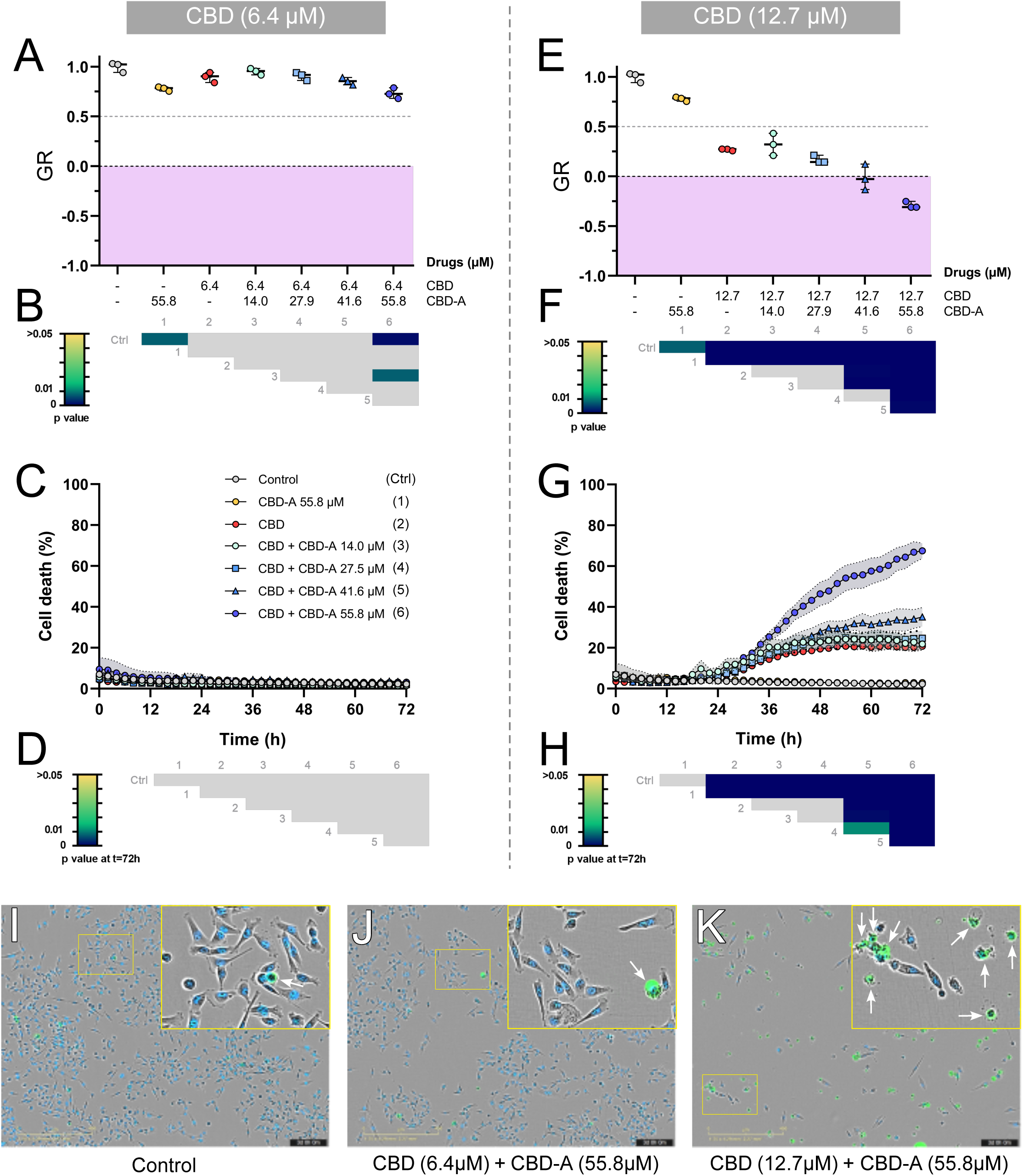
Validation of potential antagonism between CBD and CBD-A. Growth rate (GR) values for controls (DMSO 0.5%) and CBD + CBD-A combinations: CBD at 6.4 µM combined with four doses of CBD-A **(A-B)**, and CBD at 12.7 µM combined with four doses of CBD-A **(E-F)**. The regions delineated in pink in panels A and E correspond to the cytotoxic GR values. Cell death (%) for controls (DMSO 0.5%) and CBD + CBD-A combinations: CBD at 6.4 µM combined with four doses of CBD-A **(C-D)**, and CBD at 12.7 µM combined with four doses of CBD-A **(G-H)**. Data are presented as means ± SD from three independent experiments, and statistical analysis was performed using one-way ANOVA followed by Tukey’s post hoc test for multiple comparisons. A heatmap summarising the multiple comparisons is provided below each graph **(B, D, F, H).** Ctrl: control; 1 to 6 correspond to the experimental conditions in the same order as listed in the figure. Statistical significance was set at p < 0.05. Grey colour corresponds to non-significant p-values. **(I-K)** Representative composite images combining the phase contrast, green (SYTOX Green, dead cells), and NiR (SPY650-DNA, nuclei) fluorescence channels of MDA-MB-231 cells after 72h of treatment. **(I)** Control, **(J)** CBD 6.4 µM + CBD-A 55.8 µM, **(K)** CBD 12.7 µM + CBD-A 55.8 µM. Scale bars correspond to 400 µm. Insets represent zoomed images of the boxed regions. Dead cells (SPY^+^/SYTOX Green^+^ nuclei) are indicated by white arrows.

To comprehensively assess this polypharmacological synergy, the evaluation was expanded beyond binary combinations to include a three-drug formulation combining CBD, CBG, and BCP. The heatmaps in **Fig. 4A** display the percentage of growth inhibition for all tested combinations, with each heatmap representing a different dose of BCP combined with four doses of CBD and four doses of CBG. This representation highlights different response clusters, with combinations lacking CBD (CBD = 0) showing limited efficacy, whereas those including intermediate CBD concentrations (≈9.5 µM) and high BCP doses (≥48.9 µM) exhibited the strongest inhibitory effects. The corresponding synergy maps (**Fig. 4B**) revealed that these same combinations generated consistently positive synergy scores exceeding +50 across Bliss, ZIP, and HSA models, in the top three formulations (**Table 1**). In contrast, the Loewe model yielded lower scores. This discrepancy is likely due to the model’s strict assumption of dose-additivity, which is primarily designed for compounds with similar modes of action. Thus, Loewe may be less representative for multi-drug systems targeting divergent signalling nodes [23]. The three selected top synergistic combinations achieved over 70% growth inhibition in the 72 h metabolic endpoint assay. Extended data on synergy scores, combination index, and percentage inhibition for all tested combinations are provided in **Table S4**.

**Figure 4:**
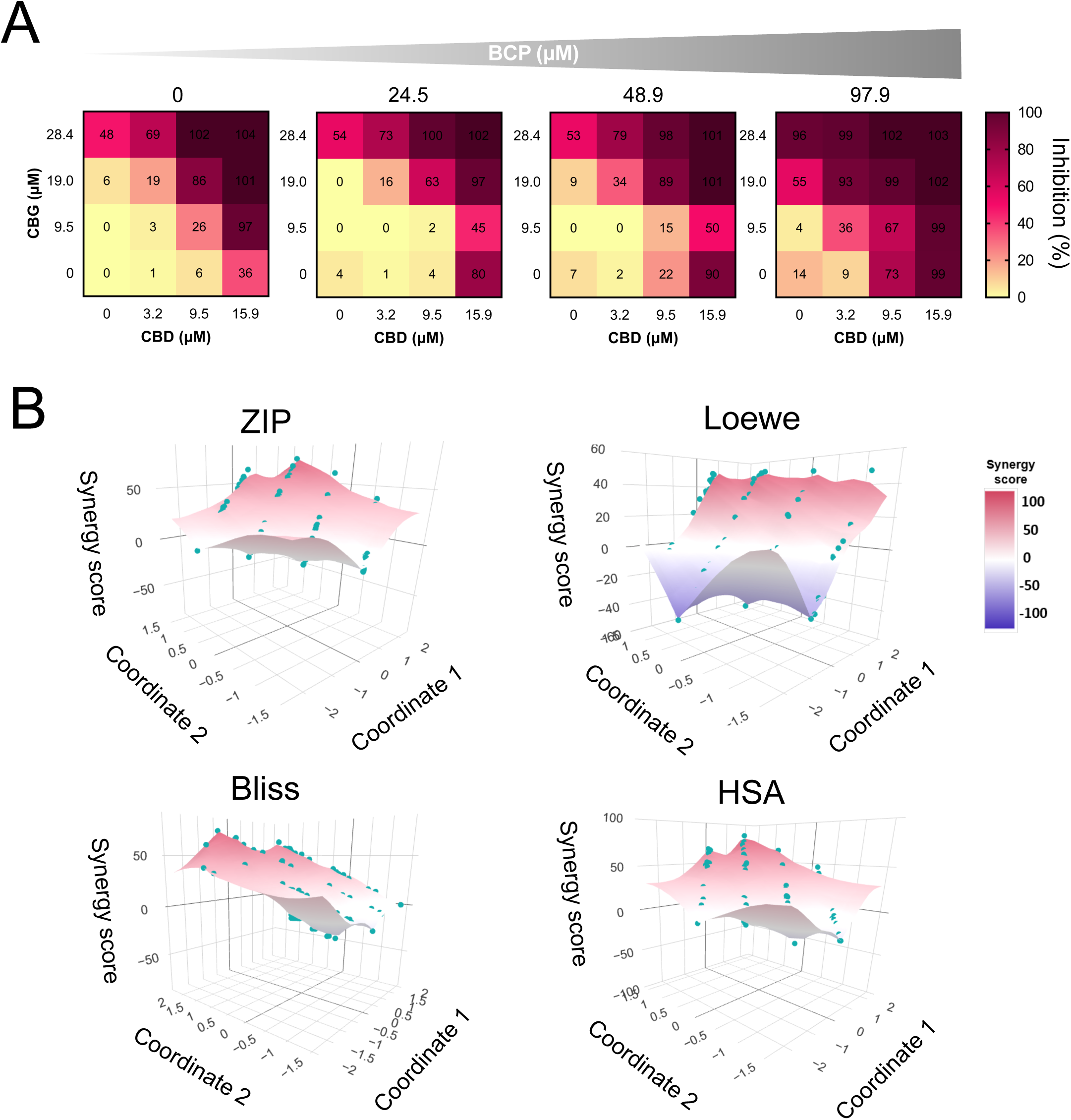
Dose-response matrix and synergy landscape analysis for the CBD, CBG and BCP combinations. **(A)** Dose-response heatmaps displaying the percentage of growth inhibition across the dose matrix of CBD and CBG, combined with BCP at 0, 24.5, 48.9, and 97.9 µM in MDA-MB-231 cells after 72 h of treatment, as determined by resazurin-based metabolic assay. **(B)** Three-dimensional synergy-score landscape generated using four reference models (ZIP, Loewe, Bliss, and HSA). Each point corresponds to a unique dose combination, positioned using multidimensional scaling (MDS) coordinates derived from dose-ranking vectors. Colours indicating the degree of interaction: synergy (red/pink), additivity (white), or antagonism (purple). Data represent the median of three independent experiments performed in triplicate.

**Table 1.** Synergy scores and corresponding growth inhibition (%) of the three most synergistic three-drug combinations across the four reference models. Growth inhibition of MDA-MB-231 cells treated with the drug combinations (C1-3) was measured after 72 h of treatment using the vitality assay. Data are means ± SD from three independent experiments. Colour scale consistent with Fig. 4.

| n° | Drug doses (μM)<br>in combination | | | Synergy scores | | | | Inhibition % |
| --- | --- | --- | --- | --- | --- | --- | --- | --- |
|  | CBD | CBG | BCP | ZIP | HSA | Loewe | Bliss | Metabolic assay |
| **C1** | 9.5 | 19.0 | 48.9 | 67.7 | 77.4 | 34.2 | 67.7 | **84.4 ± 8.4** |
| **C2** | 9.5 | 0.0 | 97.9 | 62.6 | 66.9 | 50.6 | 61.5 | **75.4 ± 16.8** |
| **C3** | 9.5 | 19.0 | 97.9 | 75.8 | 85.7 | 47.9 | 76.2 | **99.4 ± 1.4** |

To validate these results, the cytotoxic activity of these combinations was quantified by live-cell imaging in MDA-MB-231 cells (**Fig. 5**). When tested individually, CBG (19.0 µM) and BCP (97.9 µM) showed no significant activity (GR of 0.94 and 0.83, respectively), while CBD (9.5 µM) induced a modest reduction in cell growth (GR = 0.70). In sharp contrast, the combinations were significantly more effective than single agents used at the same doses. The three most synergistic combinations produced cytotoxic effects, with mean GR values of –0.25 and –0.55, for C1 and C3, respectively and a cytostatic effect (GR = 0.13) for C2 (**Fig. 5A**). All these combinations included CBD at 9.5 µM, underscoring the need for a minimal CBD concentration to maintain a cytostatic/cytotoxic activity. Each combination involved sub-active doses, with about half of the IC₅₀ of CBD, two-thirds for CBG, and up to two-thirds for BCP, supporting the dose-reduction benefit of combination therapy.

**Figure 5.**
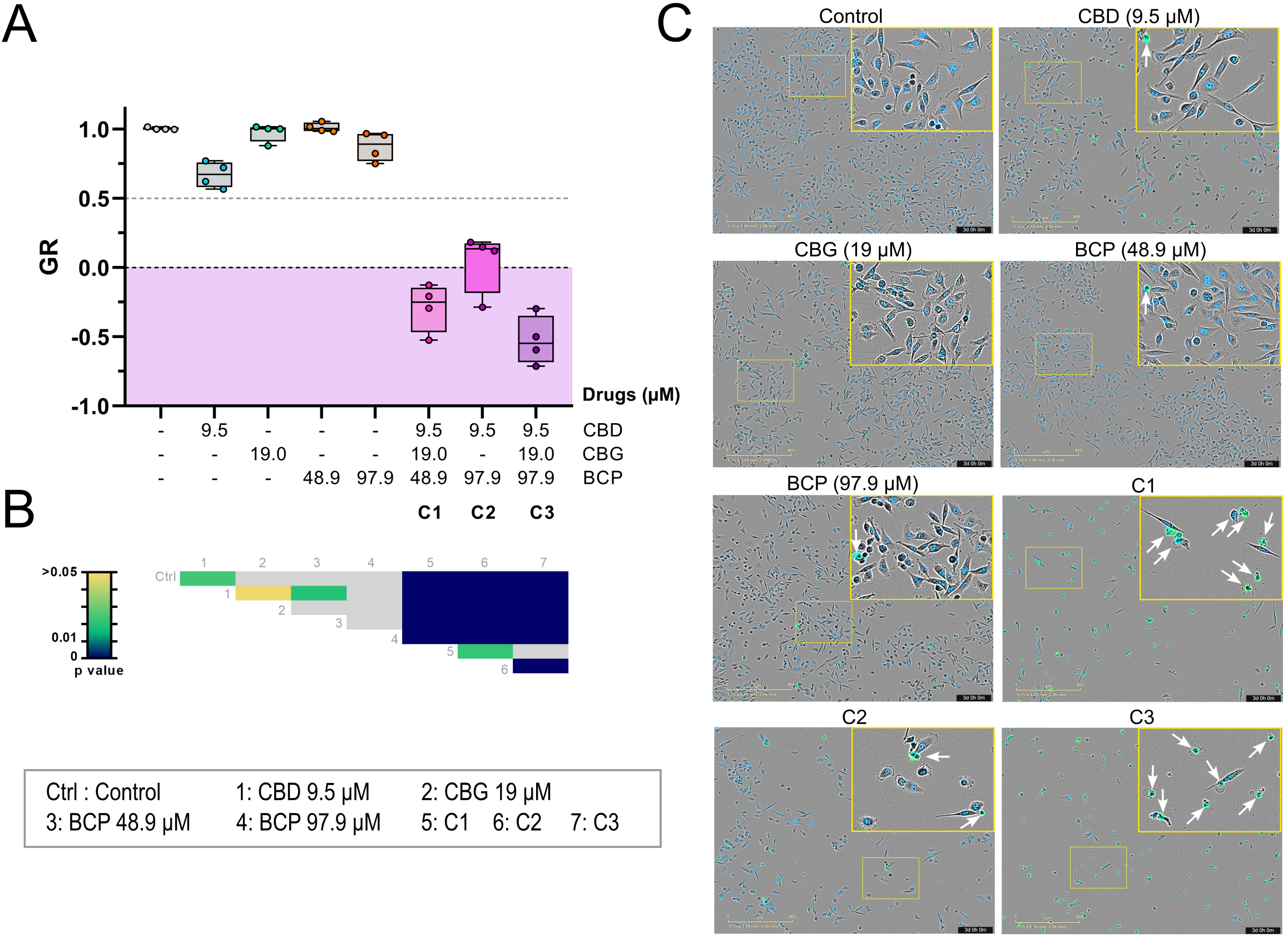
Analysis of selected three-drug combinations by live-cell imaging in MDA-MB-231 cells. MDA-MB-231 cells were treated during 72h with the three most potent combinations of CBD, CBG and BCP (C1-C3) or their individual components. DMSO (0.3%) was used as the vehicle control. **(A)** Growth rate (GR) values were calculated for each combination. Data represents the median of four independent experiments and were analysed using one-way ANOVA followed by Tukey’s post hoc test for multiple comparisons. A heatmap summarising the multiple comparisons is provided below the graph **(B).** Ctrl: control; 1 to 7 correspond to the experimental conditions in the same order as listed in the figure. Statistical significance was set at p < 0.05. Grey colour corresponds to non-significant p-values. **(B) (C)** Representative composite images combining the phase contrast, green (SYTOX Green, dead cells), and NiR (SPY650-DNA, nuclei) fluorescence channels of MDA-MB-231 cells after 72h of treatment. Scale bars correspond to 400 µm. Insets represent zoomed images of the boxed regions. Dead cells (SPY^+^/SYTOX Green^+^ nuclei) are indicated by white arrows.

Despite these reduced doses, the C1 and C3 combinations strongly induced cell death, as demonstrated by a reduced cell density associated with the presence of a high number of SYTOX Green+ dead cells compared to the control and single agents. In accordance with its cytostatic activity, C2 resulted in a decrease in cell density, accompanied by a modest increase in cell death (**Fig. 5C**).

### 3.3. Kinetic analysis of the top three combinations on MDA-MB-231 cell viability

To further characterise the biological activities of these combinations, live-cell analyses were performed to generate kinetic data (**Supplementary videos S1-S4**). Each combination was compared with its corresponding single agents to evaluate the benefit of combination therapy. **Figures 6A-C** show the time-dependent evolution of the GR values, reflecting the dynamic effect of each treatment on the population of cancer cells. C1 and C3 induced an early decline in cell growth within the first 36h of treatment, indicating a rapid onset of cytotoxicity. In contrast, C2 maintained GR values close to 0, consistent with a cytostatic effect. Crucially, all three combinations were significantly more effective than their respective single agents. Among the individual agents, only CBD (9.5 µM) was able to induce a measurable growth inhibition. Given that overall cell growth is defined by the relative balance between proliferation and death, we individually quantified both parameters. Quantification of cell proliferation was based on the evolution of the number of SPY^+^ nuclei (**Fig. 6D-F**). This analysis confirmed a significant antiproliferative activity for all three combinations. Specifically, C1 and C3 induced a net decrease in cell number compared to the initial count (*t*=0).

**Figure 6:**
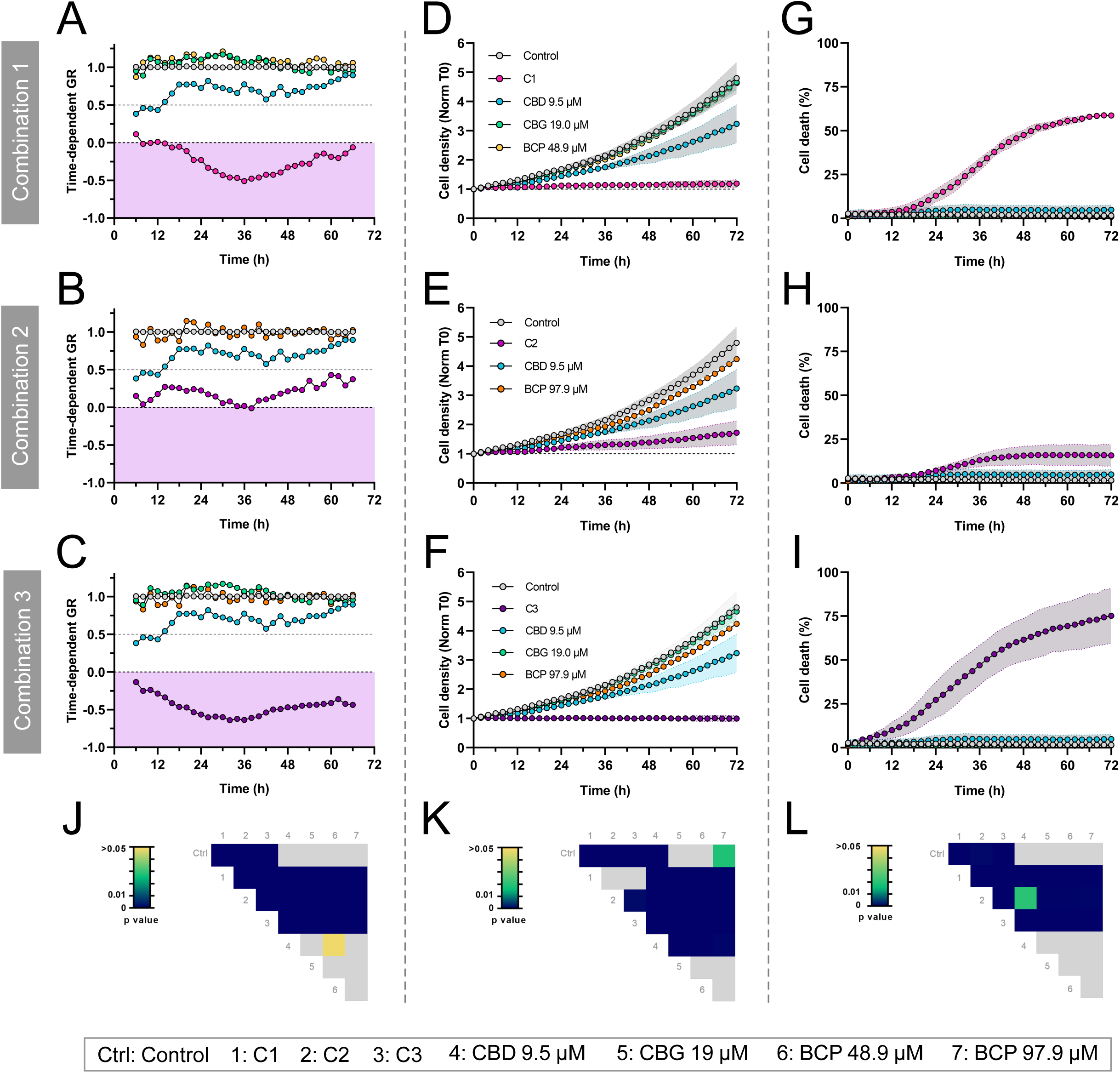
Kinetic analysis of the top synergistic combinations on the growth of MDA-MB-231 cells. MDA-MB-231 cells were treated with vehicle control (0.3% DMSO) or with combinations C1, C2 or C3 as described in **Table 1**. Cell responses were analysed by live-cell imaging for 72h, and automated image analysis was performed to quantify the time-dependent GR values **(A-C)**, the evolution of cell density **(D-F)** and cell death **(G-I)**. For each combination, the corresponding control and single-agent treatments are shown on the same graph. the time-dependent GR values following exposure to the different treatments. **(A-C)** Time-dependent GR values were estimated over 6-hour intervals. The region delineated in pink in the figure corresponds to the cytotoxic GR values. **(D-F)** Cell growth was quantified by measuring the density of SPY-DNA^+^ nuclei, normalised to the initial density at T0. **(G-I)** The percentage of dead cells was defined as the proportion of SYTOX Green^+^ nuclei normalised to SPY-DNA^+^ nuclei. Data are expressed as mean + SD from three independent experiments (in triplicate). Statistical analyses were performed using one-way ANOVA followed by Tukey’s post hoc test for multiple comparisons. A heatmap summarising the multiple comparisons is provided below each graph. **(J)** statistical analysis on the GR value after 72h of treatment. **(K-L)** Statistical analysis was performed on the area under the curve (AUC) of each individual curve. Statistical significance was set at p < 0.05. Grey colour corresponds to non-significant p-values.

In contrast, the corresponding single agents did not significantly alter cell proliferation compared to vehicle-treated cells, except for CBD, which induced a limited decrease in cell number. Kinetic analysis of cell death (SYTOX Green+ nuclei) revealed distinct effects among the combinations (**Fig. 6G-I**). While C1 and C3 were potent inducers of cell death, C2 induced a more modest but statistically significant effect, likely due to the absence of CBG in this formulation. Consistently, none of the single agents was cytotoxic.

To validate our findings beyond the MDA-MB-231 model, we evaluated these agents in Hs578T cells, another mesenchymal stem-like TNBC line 1 [24]. Results were largely consistent across both models, with the exception that C1 demonstrated a slightly reduced activity in Hs578T cells compared to MDA-MB-231 (**Supplementary Fig. S1-2**).

To evaluate the kinetic impact of synergistic combinations on cell population dynamics, we performed live-cell imaging over 72 hours (**Fig. 7A** and **Supplemental Videos S1–S4**). While vehicle-treated MDA-MB-231 cells exhibited robust migration and proliferation, all tested combinations significantly impaired these parameters, with a noticeable onset of cell death occurring approximately 18 hours post-treatment. Consistent with our synergy scores, combinations C1 and C3 were the most potent inducers of cell death. Detailed kinetic monitoring revealed that the cell death induced by C1 and C3 was preceded by distinct morphological alterations, including cellular retraction and the formation of intracytoplasmic vacuoles (**Fig. 7A**, blue insets), followed by extensive membrane blebbing (**Fig. 7A**, yellow insets) suggestive of an apoptotic cell death. At later stages, treated cells exhibited positive SYTOX staining and large cytoplasmic blisters, hallmarks of secondary necrosis (**Fig. 7A**, orange insets). To further characterize the death modality, co-incubation with a fluorogenic caspase-3/7 substrate (NucView) confirmed that cellular shrinkage and caspase activation preceded the terminal loss of membrane integrity (**Fig. 7B**), verifying an initial apoptotic execution phase. However, the limited resolution of the current imaging technique was insufficient to permit a detailed morphological and mechanistic analysis of the cellular processes induced by these treatments.

**Figure 7:**
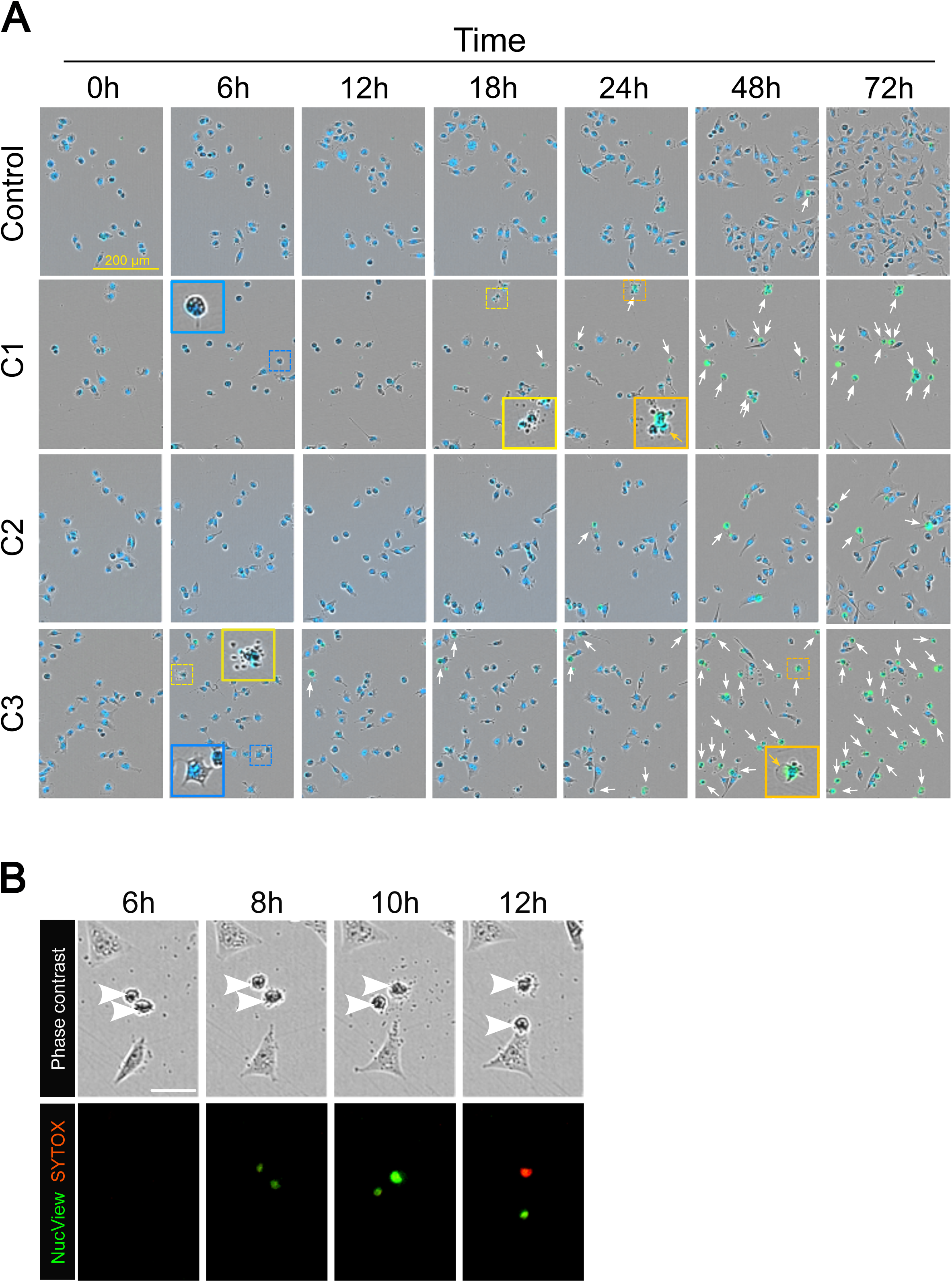
Live-cell imaging reveals morphodynamic alterations induced by synergistic combinations. **(A)** MDA-MB-231 cells were treated with vehicle (0.3% DMSO - Control) or combinations C1 to C3 as described in **Table 1** in the presence of SYTOX Green and SPY-DNA. Automated live-cell imaging was performed for 72h and representative composite images combining the phase contrast, green (SYTOX Green, dead cells), and NiR (SPY650-DNA, nuclei) fluorescence channels obtained at different time points are displayed. Dead cells (SPY^+^/SYTOX Green^+^ nuclei) are indicated by white arrows. Insets show higher magnifications of specific elements. Blue boxes highlight retracted cells exhibiting cytoplasmic vacuoles, yellow boxes indicate cells displaying extensive cytoplasmic blebbing, and orange boxes highlight the formation of large cytoplasmic blisters (orange arrows) characteristic of secondary necrosis. Scale bar corresponds to 200 µm. See **Videos S1-4** (Supplementary Material) for the complete 72 h time-lapse sequences. **(B)** MDA-MB-231 cells were treated with combinations C1 in the presence of SYTOX Orange and NucView 488 Caspase-3 substrate. Automated live-cell imaging was performed and representative phase contrast and composite images combining the green (NucView, active caspase-3) and orange (SYTOX Orange, dead cells) fluorescence channels obtained at different time points are displayed. Retracted cells are indicated by white arrow heads.

### 3.4. Spatiotemporal characterisation of C1 and C3-induced cell death: a sequential transition from autosis to apoptosis

To gain subcellular resolution of the cellular response induced by the most active combinations (C1 and C3), we employed a high spatial (200 nm) and temporal (20 min) resolution label-free live cell imaging technology called holotomography to monitor phenotypic dynamics in MDA-MB-231 cells (**Fig. 8A** and **Supplemental Videos S5–S7**). In control conditions, most cells maintained a polarised morphology characterised by an extended lamellipodium at the migration front (**Fig. 8A**, Control at 2h, *yellow inset*). Active cellular migration and proliferation were observed, resulting in the establishment of confluency by the end of the 24-hour observation period. Consistent with previous observations (**Fig. 6D, F**), C1 and C3 treatment effectively abrogated MDA-MB-231 cell proliferation (**Fig. 8B**). Within the first 8 hours of exposure, cells underwent extensive morphological remodelling of the plasma membrane, cytoplasm, and nucleus. This initial phase was characterized by the retraction of extensive lamellipodia into dynamic, blebbing protrusions and ubiquitous cytoplasmic vacuolization (**Fig. 8A**). A defining feature was the focal delamination of the inner and outer nuclear membranes, leading to significant expansion of the perinuclear space (PNS) and subsequent nuclear compression (**Fig. 8A**, orange insets), changes consistent with reported autosis hallmarks [25]. Following this autotic phase, most cells transitioned into a canonical apoptotic phenotype, marked by global cell rounding, cytoplasmic shrinkage, and the extension of filamentous apoptopodia (**Fig. 8A**, blue insets) [26]. This trajectory culminated in a total loss of structural integrity and the extrusion of large cytoplasmic blisters indicative of terminal secondary necrosis [27]. While the morphodynamic sequence was largely conserved between treatments, C3 uniquely caused the early formation of larger cytoplasmic vacuoles in a subset of the cell population (**Fig. 8A**, C3 at 2h, orange inset). Quantitative time-lapse analysis revealed that C1 and C3 induced a progressive increase in dry mass density (DMD), while the control population remained stable (**Fig. 8C**). By 26 hours, DMD was significantly higher in treated groups (C1 > C3) relative to controls. This densification likely reflects the intracellular compaction driven by lamellipodia retraction and cytoplasmic condensation (**Fig. 9A**). This phenomenon, termed apoptotic volume decrease [28], is a fundamental prerequisite for the execution phase of apoptosis, mediated by the coordinated activation of specific ion channels and transporters [28, 29]. Furthermore, C1 and C3 treatment triggered an abrupt decline in cell granularity during the first 10 hours (**Fig. 8D**), mirroring the rapid accumulation of cytoplasmic vacuoles and PNS expansion. Simultaneously, a steady decrease in cell eccentricity was observed (**Fig. 8E**), quantifying the progressive cell rounding as cells retracted membrane protrusions and underwent apoptotic shrinkage (**Fig. 9A**). This high-frequency time-lapse imaging enabled the precise delineation of cell death kinetics and single-cell penetrance, mapping the phenotypic trajectory from initial subcellular remodelling to terminal lysis (**Fig. 9A, B** and **Supplementary Fig. S3**). C1 treatment induced widespread cytoplasmic vacuolization in most cells (72/78 cells; 92.3%), a rapid response already detectable at the onset of imaging (2 h post-treatment). Within this vacuolated population, 87.5% (63/72) progressed to the distinctive PNS swelling that defines the autotic pathway [25, 30]. Notably, in a subset of cells (9/78; 11.5%), PNS swelling was the primary morphological alteration detected, preceding or bypassing visible vacuole formation and suggesting an accelerated autotic entry. In total, 92.3% (72/78) of the tracked population exhibited PNS swelling. While highly penetrant, the onset of this phenotype was markedly heterogeneous, with a median initiation time of 6.0 h post-treatment (IQR = 5.3–7.0 h). Among autotic cells, 68% (49/72) subsequently transitioned to an apoptotic execution phase, defined by cytoplasmic shrinkage followed by plasma membrane blebbing [31]. The transition from autotic PNS swelling to apoptotic blebbing exhibited a median lag time of 11.3 h and was characterized by profound kinetic heterogeneity (IQR = 7.7–16.8 h). Furthermore, 22.2% (16/72) of autotic cells progressed to apoptotic shrinkage but did not reach the blebbing stage within the 26-h observation window. A minor subpopulation (4/72; 5.6%) bypassed apoptotic blebbing to undergo direct necrotic collapse, characterized by the extrusion of large cytoplasmic blisters. Only one cell (1.3%) reached apoptosis without prior PNS swelling, and a single cell remained morphologically normal throughout the experiment. Collectively, this single-cell mapping identifies a dominant, multimodal cell death program in C1-treated MDA-MB-231 cells, defined by a structured kinetic transition from autotic remodelling to apoptotic execution.

**Figure 8:**
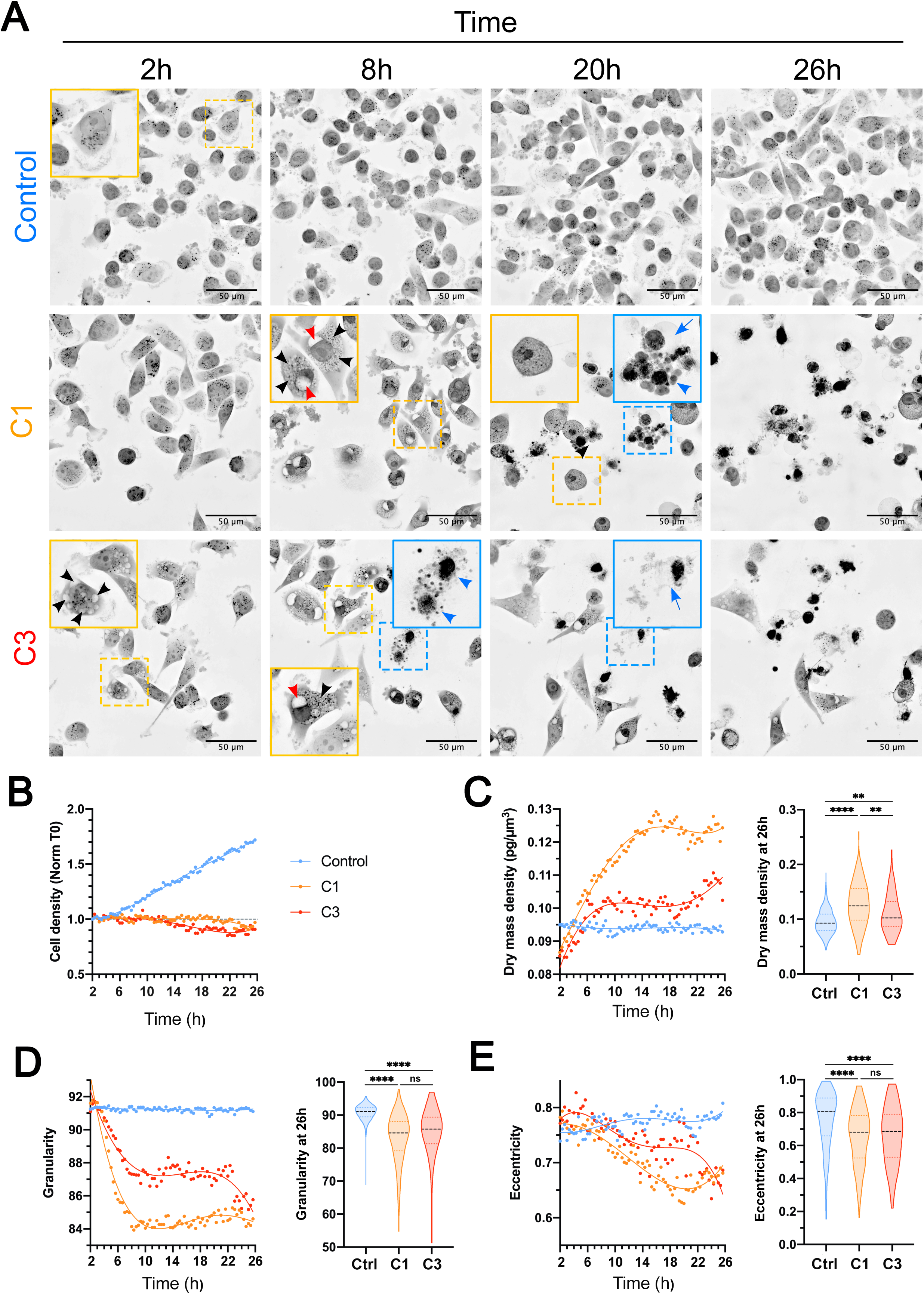
Early phenotypic alterations induced by synergistic. 932 **combinations.** MDA-MB-231 cells were treated with vehicle (0.3% DMSO - Control) or synergistic combinations C1 and C3 and monitored via label-free live-cell holotomographic microscopy (HTM). **(A)** Kinetic monitoring started 2 hours post-treatment and continued for 24 h. Images represent 2D maximum intensity projections of the Refractive Index (RI) along the Z-axis, displayed using an inverted lookup table where grey levels correlate with the RI of the samples. Insets show higher magnifications of subcellular features. Red, black, and blue arrowheads denote perinuclear swelling, intracytoplasmic vacuoles, and apoptotic-like cell fragmentation, respectively. Blue arrows indicate features of secondary necrosis. Scale bars: 50 μm. Representative timepoints at 2, 8, 20, and 26 h post treatment are shown (see **Videos S5–S7** for full sequences). **(B–E)** Quantitative HTM-based morphometric analysis performed with Eve Explorer. Temporal evolution of **(B)** cell density, **(C)** median dry mass density, **(D)** median cell granularity, and **(E)** median cell eccentricity. Violin plots illustrate single-cell distributions at 26 h post treatment; horizontal dashed lines represent the median. N = 404 (Control), 163 (C1), and 79 (C3). Statistical significance was determined via Kruskal-Wallis test followed by Dunn’s post hoc test; **** *p* < 0.0001; ** *p* < 0.001; ns: not significant.

**Figure 9:**
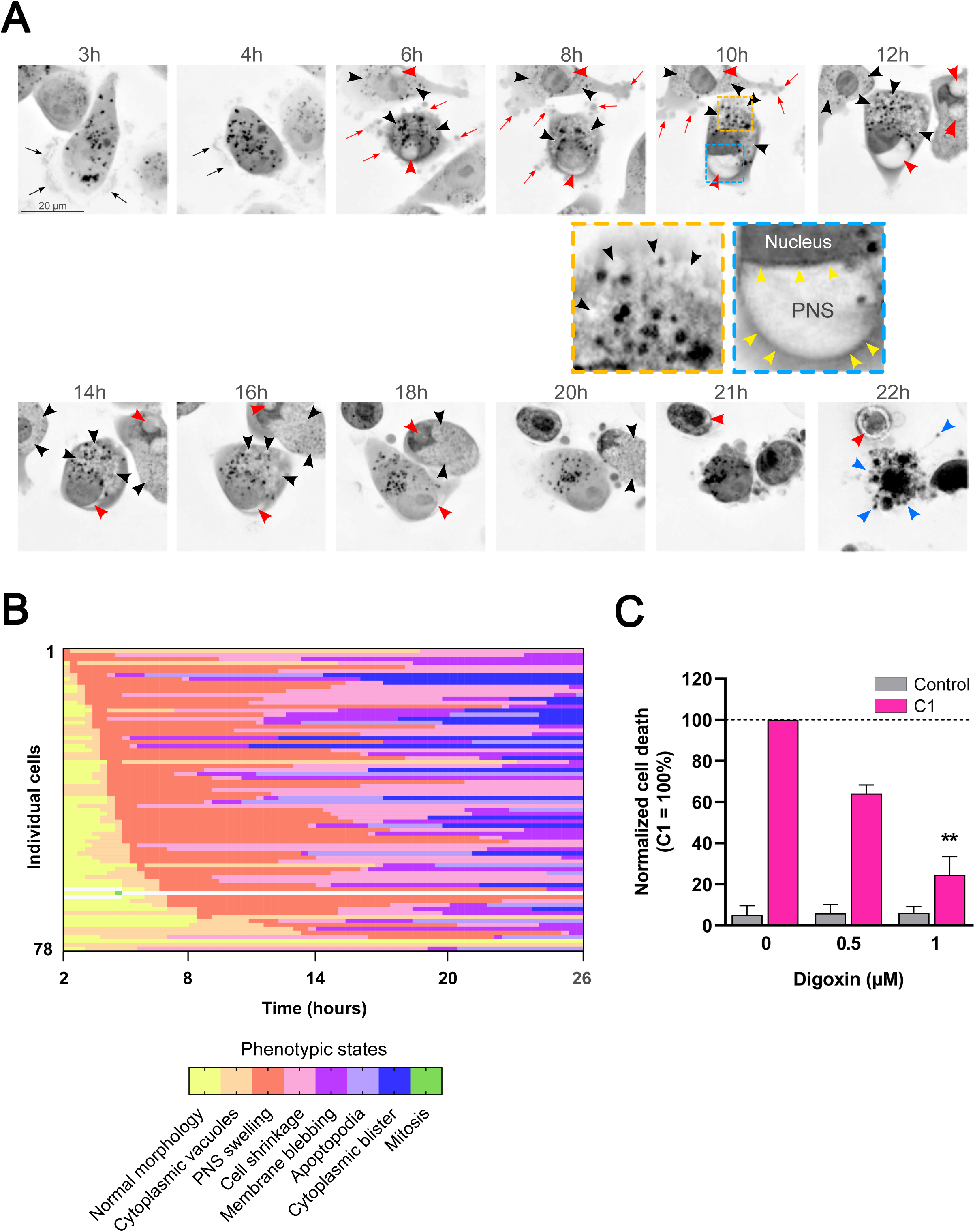
Synergistic C1 treatment triggers an autosis-to-apoptosis switch. MDA-MB-231 cells were monitored via label-free HTM for 24 h, beginning 2 h post-C1 treatment to resolve the subcellular kinetics of cell death. **(A)** Representative 2D maximum intensity projections of the refractive index. Insets demonstrate higher magnifications of intracytoplasmic vacuoles (orange square) and focal swelling of the perinuclear space (PNS) leading to nuclear compression (blue square), hallmarks of autosis. Red, yellow, black, and blue arrowheads indicate PNS swelling, nuclear membranes, cytoplasmic vacuoles, and apoptotic fragmentation, respectively. Black and red arrows denote lamellipodia and bleb-rich protrusions. Scale bars: 20 µm. **(B)** Single-cell phenotypic trajectory analysis (fate mapping) of 78 individual C1-treated cells over 24 h. Horizontal lines represent the temporal evolution of individual cells; colours denote discrete phenotypic states. Trajectories are ordered by the onset of PNS swelling to demonstrate the kinetic conservation of the autotic phase across the population. **(C)** Pharmacological rescue of C1-induced autosis by digoxin. MDA-MB-231 cells were pre-treated for 2 h with digoxin (0.5 and 1 µM) prior to C1 exposure. Data represent relative cell death normalized to the C1-only group (100% reference baseline, dashed line) at t = 6 h. Results are presented as mean ± SD (n=3). Statistical significance was assessed by a one-sample t-test (relative to the 100% baseline); ** p < 0.01.

C3 exhibited lower penetrance and slower initiation of the autotic program, inducing PNS swelling in 82.4% of tracked cells with a delayed median onset of 7.0 h post-treatment (IQR = 5.7–8.7 h) (**Supplementary Fig. S3B** and **S3C)**. Furthermore, C3 demonstrated a lower transition rate from autotic commitment to apoptotic execution, with 50% of cells progressing to membrane blebbing (**Supplementary Fig. S3B** and **S3E**). Notably, the kinetics of this transition were similar (**Supplementary Fig. S3D)** with a median lag time of 11.3 h between the onset of PNS swelling and the initiation of membrane blebbing for both combinations, despite the differences in initiation speed and penetrance. This invariance identifies the 11.3-hour window as a conserved, drug-independent execution phase. Both combinations also induced a near-complete block of the cell cycle, with mitotic frequencies restricted to 1.3% (C1) and 3.9% (C3). A minor but consistent subpopulation in both groups (5.6% for C1 and 4.8% for C3) bypassed the apoptotic phase to undergo direct necrotic collapse via cytoplasmic blistering. Together, these comparative data identify C1 and C3 as potent inducers of the autotic-to-apoptotic program while confirming that the execution trajectory follows a conserved temporal constraint once the threshold for nuclear compromise is crossed.

While PNS swelling is a distinctive hallmark of autosis, morphological characterization alone is insufficient for definitive classification. Autosis is specifically mediated by the Na^+^,K^+^-ATPase pump, the activity of which is required for the development of its associated morphological alterations [25,32]. To mechanistically confirm the involvement of this pathway, we conducted pharmacological rescue experiments using the cardiac glycoside digoxin. Pre-treatment with digoxin (2 h) inhibited C1-induced lethality in a dose-dependent manner. This analysis was performed at 6 h post-treatment, a time point at which digoxin alone exhibited no detectable cytotoxicity (**Supplementary Fig. S4**). Although 0.5 µM digoxin resulted in a moderate but non-significant trend toward reduced lethality (64.2% relative death), 1 µM digoxin significantly attenuated the response, reducing relative cell death to 24.7% (**Fig. 9C**). This robust pharmacological rescue validates the Na^+^,K^+^-ATPase-dependent autotic program as the primary driver of C1-induced cell death in this model.

Collectively, these data demonstrate the potent, early-onset cytotoxicity of the CBD, CBG, and BCP combination, which induces a previously uncharacterized cell death modality, initiating with the hallmark features of autosis before progressing to apoptosis.

## 4. Discussion

The present study addresses an unmet pharmacological need in TNBC, a subtype characterised by the absence of targetable hormone receptors and the overexpression of EGFR, which collectively limits therapeutic options and drives poor clinical outcomes. Using MDA-MB-231 cells as an established model of highly metastatic TNBC, we systematically characterised the cytotoxic and anti-proliferative activity of three phytocannabinoids (CBD, CBG, and CBD-A) and the sesquiterpene β-caryophyllene (BCP), both as single agents and in combination.

The antiproliferative activity of CBD in MDA-MB-231 cells is consistent with prior reports implicating the EGF/EGFR axis as a relevant target in this context [33] and extends earlier work by providing GR-normalised potency data suitable for quantitative combination analysis. CBG, by contrast, remains poorly characterised in breast cancer models; the present data provide quantitative evidence of its antiproliferative and cytotoxic potential, adding to the limited pharmacological profile of this cannabinoid in oncology. CBD-A exhibited modest cytotoxicity only at the highest concentrations tested and was excluded from combination studies, both for its limited efficacy and its well-documented chemical instability: thermal decarboxylation to CBD during standard extraction and handling procedures compromises reproducibility. Although CBD-A may retain pharmacological relevance in other contexts, including anti-migratory activity in MDA-MB-231 cells [36], its physicochemical liabilities preclude its inclusion in a rigorous combination design.

Three binary and ternary combinations of CBD, CBG, and BCP emerged as the most potent growth-inhibiting treatments. Because conventional IC₅₀-based metrics do not account for differences in cell division rates across experimental conditions, drug responses were quantified using growth rate (GR) inhibition metrics, which normalise drug effect to the proliferation rate of untreated controls and thus provide a more informative measure of compound potency [17]. A critical observation, only accessible through GR-based quantification, was that these three combinations produced fundamentally different biological outcomes despite appearing equally effective in endpoint metabolic assays. Combinations C1 and C3 drove net cytotoxicity (GR < 0), whereas C2 produced predominantly cytostatic effects (GR near 0). This distinction has direct therapeutic implications: cytotoxic combinations eliminate cells, while cytostatic ones arrest their proliferation without reducing viable cell number, a difference that is masked entirely by endpoint viability readouts calibrated at a single time point.

Synergy quantification was performed using SynergyFinder+, which integrates four reference models (HSA, Bliss, ZIP, and Loewe) across full dose-response matrices [19]. Combinations lacking CBD consistently showed reduced efficacy and lower synergy scores across all models, confirming CBD as the indispensable pharmacological anchor of the active combinations. Among the four models, Loewe additivity consistently yielded the lowest synergy scores, an outcome that reflects a conceptual limitation of this model in the present context. Loewe additivity assumes mutual dose-substitutability between agents, which is only valid when compounds share identical mechanisms of action. CBD, CBG, and BCP engage partially overlapping but pharmacologically distinct targets: both CBD and CBG modulate TRPV1 channels and PPARγ signalling, while BCP acts as a selective CB₂ receptor agonist [34–35]. Given this partial, rather than complete, target overlap, Bliss independence and the related ZIP model, both of which are grounded in the assumption of probabilistic independence between agents, provide a more appropriate framework for interpreting multi-drug interactions in this setting [19].

The partial additivity observed for CBD and CBG combinations aligns mechanistically with their shared engagement of TRPV1 and PPARγ: when two compounds modulate the same pathway, their effects tend toward reinforcement rather than synergistic amplification. In contrast, CBD–BCP combinations displayed marked synergy, consistent with the non-redundant character of their respective mechanisms. BCP, as a selective CB₂ agonist, inhibits cancer cell proliferation through a pathway that is largely distinct from CBD’s multi-target pro-apoptotic activity, which operates through PPARγ upregulation and the concurrent suppression of mTOR and cyclin D1 [37–39]. Although CBD has negligible intrinsic affinity for CB₂, its combination with a selective CB₂ agonist enables concurrent engagement of CB₂-dependent and CB₂-independent death pathways, a pharmacological complementarity consistent with previously reported synergistic interactions between these two compounds in analgesic models [40].

At the cellular level, high-resolution label-free holotomographic imaging provided a morphodynamic characterisation of the death process triggered by the most efficacious combinations. Combinations containing BCP (C1 and C3) induced extensive cytoplasmic vacuolisation as an early and prominent feature, consistent with the effects reported for THC, CBD, and CBG on breast cancer cell lines, where vacuolisation has been attributed to ER stress and dysregulation of autophagic flux [13, 41]. The subsequent appearance of caspase-3 activation, cell shrinkage, cytoplasmic blebbing, and apoptopodia confirmed a terminal apoptotic phase, consistent with the pro-apoptotic profiles established for CBD [13], CBG [42] and BCP [43]. However, the ordered morphological sequence observed here: vacuolisation associated with focal swelling of the perinuclear space preceding plasma membrane blebbing, is not consistent with canonical apoptosis. In contrast, the morphological intermediary phase, characterised by focal swelling of the perinuclear space and progressive nuclear compression occurring before membrane fragmentation, corresponds to the hallmarks of autosis, a regulated form of cell death mechanistically distinct from apoptosis, necroptosis, and ferroptosis. Despite its emerging recognition in several pathological contexts, autosis remains poorly characterised in breast cancer, and only one prior study has documented its occurrence in TNBC specifically. Zhang et al. demonstrated that a Beclin 1-targeting peptide, Tat-SP4, triggered autotic cell death in MDA-MB-231 and two additional TNBC cell lines, exploiting an intrinsic vulnerability rooted in the haploinsufficiency of Beclin 1, which is monoallelically deleted in 40–75% of breast cancers and is expressed at particularly low mRNA levels in TNBC relative to other breast cancer subtypes [32]. The resulting autophagy deficiency, rather than conferring resistance, paradoxically renders TNBC cells susceptible to autosis: when residual autophagic capacity is overwhelmed, even a moderate pharmacological increase in autophagy induction can exceed the cell’s homeostatic threshold and precipitate autotic death [32]. The present findings extend this paradigm to a pharmacologically distinct class of inducers, demonstrating that phytocannabinoid–terpene combinations engage the same vulnerability through a completely different mechanistic entry point (without direct Beclin 1 targeting) and through entirely distinct upstream receptor pharmacology.

Autosis is defined by two absolute requirements: (i) functional autophagy machinery, since blockade of Beclin 1, ATG5, ATG7, ATG13, or ATG14 abolishes the phenotype, and (ii) activity of the Na⁺, K⁺-ATPase pump [25]. Its biochemical definition is further refined by its resistance to caspase inhibitors, necroptosis inhibitors, and ferroptosis inhibitors, which collectively exclude it from all major previously defined regulated death pathways. The morphological sequence of autosis follows a staged ultrastructural programme: an early phase of autophagosome accumulation with convoluted nuclear membranes and dilated ER; a mid-phase of inner/outer nuclear membrane separation; and a late phase of focal swelling of the perinuclear space, which exerts mechanical pressure on the nucleus, inducing its characteristic concave deformation [25,30,32]. Autotic cells maintain substrate adhesion throughout, in direct contrast to the detachment that typifies apoptosis, a feature directly observable in live-cell imaging.

The molecular axis governing autosis centres on a direct physical interaction between Beclin 1 and the α-subunit of Na⁺,K⁺-ATPase. This interaction is dynamically regulated: it increases substantially under autosis-inducing conditions such as starvation and ischaemia, and is disrupted by cardiac glycosides, whose protective effect against autosis appears to operate at least partly through blocking this protein–protein interaction rather than solely through enzymatic inhibition of the pump [25].

The trigger for autosis is the net accumulation of autophagosomes, which can arise from excessive induction of autophagosome biogenesis, from a blockade of autophagosome-lysosome fusion, or, as is particularly relevant in TNBC, from an insufficient baseline lysosomal clearance capacity that is rapidly saturated upon pharmacological stimulation. In the context of cannabinoid action, CBD, CBG, and BCP individually induce autophagic flux in breast cancer cells [13, 41, 42, 43]; the present data suggest that at the concentrations deployed in C1 and C3, their combination generates a net surplus of autophagosomes that exceeds the degradative capacity of MDA-MB-231 cells, precipitating autotic engagement. This interpretation is fully consistent with the model established by Zhang et al., in which TNBC cells with low Beclin 1 expression tolerate basal autophagy yet lack the spare capacity to absorb a pharmacologically imposed increase in autophagic load [32].

The morphodynamic analysis further identified a temporally invariant feature of the death programme: although C1 induced a higher rate of entry into the autotic phase than C3 (92.3% vs. 82.4% of cells, respectively), the interval between perinuclear space swelling and the transition to apoptotic blebbing was fixed at a median of 11.3 hours for both combinations. This kinetic invariance across pharmacologically distinct stimuli suggests that the autosis-to-apoptosis transition is governed not by upstream induction conditions, but by an intrinsic cellular threshold, likely reflecting the time required for irreversible depletion of bioenergetic reserves, disruption of ionic homeostasis beyond a critical point, or the accumulation of sufficient organellar damage to engage caspase-dependent membrane fragmentation. The conservation of this temporal window is consistent with the classification of the autosis-to-apoptosis transition as a non-stochastic, hard-wired programme in which the upstream stimulus modulates commitment probability, but not the kinetics of execution once commitment has occurred. To the best of our knowledge, this sequential engagement of autosis followed by a transition to apoptotic execution, as a single, temporally ordered death programme, has not been previously documented in any cellular model. This point deserves emphasis: Zhang et al., in the only prior report of autosis in TNBC, described autosis as the terminal event itself, culminating in necrotic plasma membrane rupture; rescue by the Na⁺,K⁺-ATPase inhibitor digoxin was complete, and no subsequent caspase-dependent phase was identified [32]. The present data indicate a fundamentally different trajectory: perinuclear space swelling systematically precedes, and is kinetically decoupled from, subsequent apoptotic blebbing, providing evidence that autosis and apoptosis can operate as consecutive modules within a single integrated death cascade rather than as independent or parallel processes. Critically, prior studies reporting the co-occurrence of autophagic and apoptotic features in cannabinoid-treated cancer cells established co-occurrence as a morphological observation, not mechanistic succession [13, 41]. This distinction implies that the upstream autotic phase represents a discrete, pharmacologically accessible commitment point that precedes, and may be required for, the subsequent apoptotic collapse.

From a therapeutic standpoint, the present findings carry two notable implications. First, the cannabinoid–terpene combinations characterised here achieve strong cytotoxic efficacy at sub-IC₅₀ concentrations of each component, a dose-sparing effect that reduces the likelihood of off-target toxicity while potentially broadening the therapeutic window. Second, the involvement of autosis as an initiating mechanism diversifies the death pathways engaged: because autosis is independent of caspases and is not suppressed by the resistance mechanisms that typically attenuate apoptosis (e.g., Bcl-2 overexpression, p53 loss), a combination strategy that triggers autotic entry before transitioning to apoptotic execution may be less susceptible to the resistance acquisition that limits single-agent apoptosis-based approaches. The polypharmacological nature of these compounds, engaging CB₂, TRPV1, PPARγ, mTOR, and the autophagy–Na⁺,K⁺-ATPase axis concurrently, may further reduce the selective pressure driving resistance emergence. That this cytotoxic programme can be elicited in MDA-MB-231 cells from two mechanistically unrelated upstream stimuli (Beclin 1-targeting peptides [32] and phytocannabinoid–terpene combinations, as shown here) reinforces the view that autotic susceptibility is a tractable and reproducible pharmacological vulnerability in TNBC.

In summary, CBD, CBG, and BCP act synergistically in MDA-MB-231 cells to induce cytotoxicity through a temporally ordered death programme consistent with autotic initiation followed by apoptotic execution. This sequence, demonstrable only through kinetic live-cell imaging, illustrates a mechanistic complexity that is invisible to endpoint assays and underscores the value of morphodynamic characterisation in preclinical drug combination studies. Building on the single prior report of autosis in TNBC, the present data extend this emerging paradigm to phytocannabinoid–terpene combinations and reveal, for the first time, that autosis can serve as an upstream commitment module within a sequential death programme rather than as a terminal event per se. The identification of Na⁺,K⁺-ATPase-regulated autotic death as a mechanism accessible to pharmacologically diverse inducers opens a therapeutic avenue that is orthogonal to conventional apoptosis-targeted strategies in TNBC.

The use of high-purity reference compounds in the present study was a deliberate methodological choice, enabling unambiguous attribution of observed effects to defined pharmacological entities rather than to batch-dependent compositional variation. This reductionist approach provides a rational, mechanism-grounded framework directly transposable to complex formulations: hemp chemotypes naturally co-expressing CBD, CBG, and BCP, or standardised phytopharmaceutical preparations enriched in these constituents, represent immediate translational vehicles for the combinations characterised here. This perspective aligns with the observations of Blasco-Benito et al., who demonstrated that full-spectrum cannabis extracts produced markedly greater anticancer effects than purified THC alone, an outcome now interpretable considering the cooperative pharmacodynamic interactions documented in the present work [44].

## 5. Conclusion

CBD, CBG, and BCP act synergistically in TNBC cells to produce cytotoxicity at sub-IC₅₀ concentrations through a temporally ordered death programme in which autosis, a Na⁺,K⁺-ATPase-regulated, autophagy-dependent form of cell death, precedes apoptotic execution. This autosis-to-apoptosis sequence, which to the best of our knowledge has not been reported previously in any cellular model, exploits an intrinsic vulnerability of TNBC rooted in its limited spare autophagic capacity, and is accessible to phytochemical combinations acting through receptor mechanisms entirely distinct from direct Beclin 1 targeting. These findings support a polypharmacological strategy that simultaneously engages complementary, non-redundant death pathways while reducing individual compound doses, offering a rationale for the development of optimised cannabinoid-based formulations against TNBC.

## Abbreviations

BCP: β-caryophyllene
CB₂: cannabinoid receptor 2
CBD: cannabidiol
CBD-A: cannabidiolic acid
CBG: cannabigerol
DMEM: Dulbecco’s Modified Eagle Medium
DMSO: dimethyl sulfoxide
EGR: epidermal growth factor
EGFR: epidermal growth factor receptor
FBS: fetal bovine serum
GR: growth rate
HSA: Highest Single Agent
HTM: holotomographic microscopy
IC₅₀: half maximal inhibitory concentration
PPARγ: peroxisome proliferator-activated receptor gamma
THC: tetrahydrocannabinol
TNBC: triple-negative breast cancer
TRPV1: transient receptor potential vanilloid 1
ZIP: Zero Interaction Potency.

## Acknowledgments.

The authors sincerely thank all members of the Pharmacognosy and LBTD laboratories for their support. The authors thank Sandra Ormenese, Alexandre Hego and Gaetan Lefevre from the GIGA Cell Imaging Platform (ULiège, Belgium) for their help and advice.

This research was supported by the F.R.S.–FNRS (National Fund for Scientific Research, Belgium), the University of Liège (Belgium), and the Fonds Léon Fredericq. CH is funded by a FRIA doctoral fellowship from the F.R.S.–FNRS (grant number 40021906). EM is a Research Associate of the F.R.S.–FNRS.

## Supplementary Table legends

**Supplementary Table S1.** Drug concentrations (µg/mL and µM). synergy scores and combination index (CI) for CBD-CBG combination in MDA-MB-231 cells.

**Supplementary Table S2.** Drug concentrations (µg/mL and µM). synergy scores and CI for the CBD-CBD-A combination in MDA-MB-231 cells.

**Supplementary Table S3.** Drug concentrations (µg/mL and µM). synergy scores and CI for the CBD-BCP combination in MDA-MB-231 cells.

**Supplementary Table S4.** Drug concentrations (µg/mL and µM). synergy scores and CI for the three-drug combination (CBD+CBG+BCP) in MDA-MB-231 cells.

## Supplementary Figures Legends

**Supplementary Figure S1. Kinetic analysis of the top synergistic combinations on the growth of Hs578T cells.** Cell responses were analyzed by live-cell imaging for 72h, and automated image analysis was performed to quantify the time-dependent GR values (A-C), the evolution of cell density (D-F) and cell death (G-I). Data are expressed as mean + SD from three independent experiments (in triplicate). Statistical analyses were performed using one-way ANOVA followed by Tukey’s post hoc test for multiple comparisons. A heatmap summarizing the multiple comparisons is provided below each graph. (J) statistical analysis on the GR value after 72h of treatment. (K-L) Statistical analysis was performed on the area under the curve (AUC) of each individual curve. Statistical significance was set at p < 0.05. Grey colour corresponds to non-significant p-values.

**Supplementary Figure S2. Live-cell imaging reveals morphodynamic alterations induced by synergistic combinations in Hs578T cells**. Hs578T cells were treated with vehicle (0.3% DMSO - Control) or combinations C1 to C3 as described in Table 1 in the presence of SYTOX Green and SPY-DNA. **(A)** Automated live-cell imaging was performed for 72h and representative composite images combining the phase contrast, green (SYTOX Green, dead cells), and NiR (SPY650-DNA, nuclei) fluorescence channels obtained at different time points are displayed. Dead cells (SPY+/SYTOX Green+ nuclei) are indicated by white arrows. Insets show higher magnifications of specific elements. Blue boxes highlight cells exhibiting cytoplasmic vacuoles. Scale bar corresponds to 200 µm. **(B)** Growth rate (GR) values were calculated for each combination on Hs578T. Data represents the median of four independent experiments and was analyzed using one-way ANOVA followed by Tukey’s post hoc test for multiple comparisons. A heatmap summarizing the multiple comparisons is provided below the graph. Ctrl: control; 1 to 7 corresponds to the experimental conditions in the same order as listed in the figure. Statistical significance was set at p < 0.05. Grey colour corresponds to non-significant p-values.

**Supplementary Figure S3. Comparative single-cell kinetic mapping of C1- and C3-induced cell death.** MDA-MB-231 cells were monitored via label-free holotomographic microscopy at 20-minute resolution for 24 h. Imaging commenced 2 h post-treatment to resolve the subcellular phenotypic trajectories. **(A–B)** Single-cell phenotypic heatmaps for C1-treated (n=78) and C3-treated (n=55) lineages. Each horizontal track represents the temporal evolution of a single cell, with colours denoting discrete phenotypic states. Trajectories are synchronized by the onset of perinuclear space (PNS) swelling. Vertical blue dotted lines indicate the median initiation time for autotic commitment (6.0 h for C1; 7.0 h for C3). **(C)** Kinetics of autosis initiation. Kaplan-Meier plot illustrating the time-to-onset of the autotic program, defined by the first appearance of focal PNS swelling. The Y-axis represents the probability of remaining in a pre-autotic state (cells yet to exhibit PNS swelling) over time. C1-treated cells exhibited a significantly faster initiation compared to the C3 (Log-rank Mantel-Cox test, *P* < 0.05). **(D)** Temporal conservation of the execution phase. Kaplan-Meier plot showing the execution lag, defined as the interval between the onset of autotic commitment (PNS swelling) and the initiation of terminal apoptotic blebbing. The Y-axis represents the probability of autotic survival (the proportion of committed cells that have not yet transitioned to the execution phase). Both treatments showed similar conversion efficiencies (Log-rank Mantel-Cox test, *P* = 0.3056). **(E)** Comparison of the major phenotypic alterations induced by C1 and C3 treatments.

**Supplementary Figure S4. Kinetic monitoring of MDA-MB-231 cell growth under digoxin treatment.** Time-course evolution of SPY+ nuclei density was quantified using automated live-cell imaging with images acquired every 2 hours over a 24 h period. Cells were treated with vehicle (control) or digoxin (0.5 µM and 1 µM). Data represent mean ± SD (n=3). This analysis confirms that at the 6 h time point (used for C1-rescue experiments), digoxin does not induce growth inhibition or cytotoxicity compared to control cells (p > 0.05).

## Supplementary Videos

**Supplementary Videos S1-S4. Analysis of selected three-drug combinations by live-cell imaging in MDA-MB-231 cells.** MDA-MB-231 cells were treated during 72h with the three most potent combinations of CBD, CBG and BCP (C1-C3), DMSO (0.3%) was used as vehicle control. Representative videos combining the phase contrast, green (SYTOX Green, dead cells), and NiR (SPY650-DNA, nuclei) fluorescence channels of MDA-MB-231 cells during 72 hours of treatment (12 frames/day). We provided videos of control, combination 1 (C1), combination 2 (C2) and combination 3 (C3).

**Video S1**: Control treated cells

**Video S2**: C1 treated cells

**Video S3**: C2 treated cells

**Video S4:** C3 treated cells

**Videos S5-S7. Subcellular phenotypic alterations induced by synergistic combinations.** MDA-MB-231 cells were treated with vehicle (0.3% DMSO - Control) or combinations of C1 and C3. Cells were imaged by label-free live-cell holotomographic microscopy. Time-lapse acquisition started 2 hours after treatment initiation and continued for 24h (3 frames/h). Images are displayed as inverted LUT.

**Video S5**: CTRL treated cells

**Video S6**: C1 treated cells

**Video S7:** C3 treated cells

