## Supplementary Tables for "Synergistic induction of a lethal Autosis-to-Apoptosis switch by Phytocannabinoids and beta-Caryophyllene in Triple-Negative Breast Cancer Cells"

**Table S1: Drug concentrations (µg/mL and µM). synergy scores and CI for CBD-CBG combination in MDA-MB-231 cells**

Inf: infinity - /: not calculated – CI: Combination Index (CI < 1, CI=1, and CI > 1 indicate synergy, additivity and antagonism, respectively). Response % corresponds to the % of inhibition of MDA-MB-231 cell vitality after 72 hours of treatment (represented by the mean of three independent experiment).

| Drug 1 | Drug 2 | Drug 1 µM [µg/mL] | Drug 2 µM [µg/mL] | ZIP synergy score | HSA synergy score | Loewe synergy score | Bliss synergy score | CI | Response % |
|---|---|---|---|---|---|---|---|---|---|
| CBD | CBG | 0.00 [0 µg/mL] | 0.00 [0 µg/mL] | 0.00 | 0.00 | 0.00 | 0.00 | / | -2.52 |
| CBD | CBG | 0.00 [0 µg/mL] | 9.48 [3 µg/mL] | 0.00 | 0.00 | 0.00 | 0.00 | / | 5.91 |
| CBD | CBG | 0.00 [0 µg/mL] | 18.96 [6 µg/mL] | 0.00 | 0.00 | 0.00 | 0.00 | / | 51.27 |
| CBD | CBG | 0.00 [0 µg/mL] | 28.44 [9 µg/mL] | 0.00 | 0.00 | 0.00 | 0.00 | / | 92.71 |
| CBD | CBG | 0.00 [0 µg/mL] | 37.92 [12 µg/mL] | 0.00 | 0.00 | 0.00 | 0.00 | / | 98.37 |
| CBD | CBG | 0.00 [0 µg/mL] | 47.40 [15 µg/mL] | 0.00 | 0.00 | 0.00 | 0.00 | / | 5.91 |
| CBD | CBG | 3.18 [1 µg/mL] | 0.00 [0 µg/mL] | 0.00 | 0.00 | 0.00 | 0.00 | / | 1.74 |
| CBD | CBG | 3.18 [1 µg/mL] | 9.48 [3 µg/mL] | -1.49 | 2.16 | -2.32 | 4.32 | / | 2.86 |
| CBD | CBG | 3.18 [1 µg/mL] | 18.96 [6 µg/mL] | 14.30 | 19.83 | 1.97 | 19.13 | 0.99 | 25.89 |
| CBD | CBG | 3.18 [1 µg/mL] | 28.44 [9 µg/mL] | 31.90 | 33.73 | 15.37 | 33.36 | 0.86 | 82.93 |
| CBD | CBG | 3.18 [1 µg/mL] | 37.92 [12 µg/mL] | 7.47 | 6.60 | 5.75 | 6.57 | 0.53 | 100.82 |
| CBD | CBG | 3.18 [1 µg/mL] | 47.40 [15 µg/mL] | 2.15 | 4.18 | 3.04 | 4.17 | 0.44 | 102.84 |
| CBD | CBG | 6.36 [2 µg/mL] | 0.00 [0 µg/mL] | 0.00 | 0.00 | 0.00 | 0.00 | / | 3.10 |
| CBD | CBG | 6.36 [2 µg/mL] | 9.48 [3 µg/mL] | 2.63 | 8.04 | 0.87 | 10.14 | 0.97 | 10.33 |
| CBD | CBG | 6.36 [2 µg/mL] | 18.96 [6 µg/mL] | 31.58 | 27.30 | -22.19 | 24.49 | 1.14 | 35.71 |
| CBD | CBG | 6.36 [2 µg/mL] | 28.44 [9 µg/mL] | 39.68 | 39.70 | 15.77 | 38.24 | 0.85 | 91.46 |
| CBD | CBG | 6.36 [2 µg/mL] | 37.92 [12 µg/mL] | 7.62 | 6.95 | 4.89 | 6.77 | 0.50 | 101.11 |
| CBD | CBG | 6.36 [2 µg/mL] | 47.40 [15 µg/mL] | 1.67 | 2.90 | 1.53 | 2.85 | 0.49 | 101.39 |
| CBD | CBG | 7.95 [2.5 µg/mL] | 0.00 [0 µg/mL] | 0.00 | 0.00 | 0.00 | 0.00 | / | 3.52 |
| CBD | CBG | 7.95 [2.5 µg/mL] | 9.48 [3 µg/mL] | 7.51 | 6.98 | -7.34 | 9.06 | 1.10 | 9.88 |
| CBD | CBG | 7.95 [2.5 µg/mL] | 18.96 [6 µg/mL] | 56.13 | 57.93 | 0.34 | 54.46 | 0.88 | 59.97 |
| CBD | CBG | 7.95 [2.5 µg/mL] | 28.44 [9 µg/mL] | 43.10 | 47.05 | 21.47 | 45.23 | 0.69 | 98.51 |

| | | | | | | | | | |
|---|---|---|---|---|---|---|---|---|---|
| CBD | CBG | 7.95 [2.5 μg/mL] | 37.92 [12 μg/mL] | 7.28 | 5.03 | 4.35 | 4.79 | 0.86 | 99.21 |
| CBD | CBG | 7.95 [2.5 μg/mL] | 47.40 [15 μg/mL] | 0.96 | 1.53 | 0.44 | 1.47 | 0.79 | 100.07 |
| CBD | CBG | 9.54 [3 μg/mL] | 0.00 [0 μg/mL] | 0.00 | 0.00 | 0.00 | 0.00 | / | 5.93 |
| CBD | CBG | 9.54 [3 μg/mL] | 9.48 [3 μg/mL] | 20.61 | 23.61 | -0.05 | 25.65 | 1.01 | 29.05 |
| CBD | CBG | 9.54 [3 μg/mL] | 18.96 [6 μg/mL] | 68.57 | 70.15 | 5.55 | 64.87 | 0.82 | 78.48 |
| CBD | CBG | 9.54 [3 μg/mL] | 28.44 [9 μg/mL] | 43.13 | 50.50 | 16.64 | 47.61 | 0.38 | 101.32 |
| CBD | CBG | 9.54 [3 μg/mL] | 37.92 [12 μg/mL] | 8.04 | 7.87 | 4.68 | 7.51 | 0.44 | 101.29 |
| CBD | CBG | 9.54 [3 μg/mL] | 47.40 [15 μg/mL] | 2.42 | 2.24 | -1.19 | 2.15 | 0.78 | 100.75 |
| CBD | CBG | 12.72 [4 μg/mL] | 0.00 [0 μg/mL] | 0.00 | 0.00 | 0.00 | 0.00 | / | 13.22 |
| CBD | CBG | 12.72 [4 μg/mL] | 9.48 [3 μg/mL] | 23.36 | 15.75 | -27.43 | 17.63 | 1.23 | 29.39 |
| CBD | CBG | 12.72 [4 μg/mL] | 18.96 [6 μg/mL] | 70.28 | 73.02 | 10.56 | 67.72 | 0.66 | 81.12 |
| CBD | CBG | 12.72 [4 μg/mL] | 28.44 [9 μg/mL] | 38.81 | 45.72 | 15.52 | 39.25 | 0.72 | 96.08 |
| CBD | CBG | 12.72 [4 μg/mL] | 37.92 [12 μg/mL] | 5.13 | 1.86 | -1.83 | 1.03 | 0.93 | 96.23 |
| CBD | CBG | 12.72 [4 μg/mL] | 47.40 [15 μg/mL] | -0.55 | -0.37 | -4.96 | -0.56 | 0.87 | 98.25 |
| CBD | CBG | 14.31 [4.5 μg/mL] | 0.00 [0 μg/mL] | 0.00 | 0.00 | 0.00 | 0.00 | / | 23.12 |
| CBD | CBG | 14.31 [4.5 μg/mL] | 9.48 [3 μg/mL] | 32.39 | 28.48 | -12.37 | 30.14 | 1.05 | 46.59 |
| CBD | CBG | 14.31 [4.5 μg/mL] | 18.96 [6 μg/mL] | 67.96 | 72.55 | 17.60 | 67.84 | 0.57 | 96.19 |
| CBD | CBG | 14.31 [4.5 μg/mL] | 28.44 [9 μg/mL] | 36.72 | 50.74 | 11.12 | 39.48 | 0.42 | 101.07 |
| CBD | CBG | 14.31 [4.5 μg/mL] | 37.92 [12 μg/mL] | 6.40 | 6.88 | 0.98 | 5.45 | 0.66 | 101.38 |
| CBD | CBG | 14.31 [4.5 μg/mL] | 47.40 [15 μg/mL] | 1.39 | 2.33 | -2.09 | 1.98 | 0.78 | 101.31 |
| CBD | CBG | 15.9 [5 μg/mL] | 0.00 [0 μg/mL] | 0.00 | 0.00 | 0.00 | 0.00 | / | 34.84 |
| CBD | CBG | 15.9 [5 μg/mL] | 9.48 [3 μg/mL] | 49.60 | 59.26 | 24.66 | 60.65 | 0.42 | 95.59 |
| CBD | CBG | 15.9 [5 μg/mL] | 18.96 [6 μg/mL] | 59.97 | 64.11 | 19.91 | 60.22 | 0.42 | 100.48 |
| CBD | CBG | 15.9 [5 μg/mL] | 28.44 [9 μg/mL] | 31.14 | 46.52 | 12.05 | 28.82 | 0.70 | 96.71 |
| CBD | CBG | 15.9 [5 μg/mL] | 37.92 [12 μg/mL] | 5.25 | 7.61 | 1.68 | 5.40 | 0.58 | 101.34 |
| CBD | CBG | 15.9 [5 μg/mL] | 47.40 [15 μg/mL] | 1.16 | 3.29 | -1.51 | 2.79 | 0.62 | 101.70 |
| CBD | CBG | 19.08 [6 μg/mL] | 0.00 [0 μg/mL] | 0.00 | 0.00 | 0.00 | 0.00 | / | 60.09 |
| CBD | CBG | 19.08 [6 μg/mL] | 9.48 [3 μg/mL] | 43.23 | 40.61 | 20.36 | 41.54 | 0.45 | 97.68 |
| CBD | CBG | 19.08 [6 μg/mL] | 18.96 [6 μg/mL] | 41.98 | 42.30 | 17.90 | 39.70 | 0.55 | 99.65 |
| CBD | CBG | 19.08 [6 μg/mL] | 28.44 [9 μg/mL] | 21.16 | 43.67 | 15.06 | 21.82 | 0.60 | 100.72 |
| CBD | CBG | 19.08 [6 μg/mL] | 37.92 [12 μg/mL] | 3.55 | 6.33 | 1.23 | 2.76 | 0.74 | 100.38 |
| CBD | CBG | 19.08 [6 μg/mL] | 47.40 [15 μg/mL] | 0.98 | 3.30 | -1.63 | 2.39 | 0.66 | 101.45 |

**Table S2: Drug concentrations (µg/mL and µM). synergy scores and CI for the CBD-CBD-A combination in MDA-MB-231 cells**

Inf: infinity - /: not calculated – CI: Combination Index (CI < 1, CI=1, and CI > 1 indicate synergy, additivity and antagonism, respectively). Response % corresponds to the % of inhibition of MDA-MB-231 cell vitality after 72 hours of treatment (represented by the mean of three independent experiment).

| Drug 1 | Drug 2 | Drug 1 µM - [µg/mL] | Drug 2 µM - [µg/mL] | ZIP synergy score | HSA synergy score | Loewe synergy score | Bliss synergy score | CI | Response % |
|---|---|---|---|---|---|---|---|---|---|
| CBD | CBD-A | 0.00 [0 µg/mL] | 0.00 [0 µg/mL] | 0.00 | 0.00 | 0.00 | 0.00 | / | 0.00 |
| CBD | CBD-A | 0.00 [0 µg/mL] | 13.95 [5 µg/mL] | 0.00 | 0.00 | 0.00 | 0.00 | / | 3.36 |
| CBD | CBD-A | 0.00 [0 µg/mL] | 27.89 [10 µg/mL] | 0.00 | 0.00 | 0.00 | 0.00 | / | 4.88 |
| CBD | CBD-A | 0.00 [0 µg/mL] | 41.84 [15 µg/mL] | 0.00 | 0.00 | 0.00 | 0.00 | / | 7.54 |
| CBD | CBD-A | 0.00 [0 µg/mL] | 55.79 [20 µg/mL] | 0.00 | 0.00 | 0.00 | 0.00 | / | 9.40 |
| CBD | CBD-A | 0.00 [0 µg/mL] | 83.68 [30 µg/mL] | 0.00 | 0.00 | 0.00 | 0.00 | / | 30.28 |
| CBD | CBD-A | 0.00 [0 µg/mL] | 111.58 [40 µg/mL] | 0.00 | 0.00 | 0.00 | 0.00 | / | 57.47 |
| CBD | CBD-A | 3.18 [1 µg/mL] | 0.00 [0 µg/mL] | 0.00 | 0.00 | 0.00 | 0.00 | / | 1.74 |
| CBD | CBD-A | 3.18 [1 µg/mL] | 13.95 [5 µg/mL] | -2.39 | -5.49 | -20.17 | -5.83 | Inf | -1.44 |
| CBD | CBD-A | 3.18 [1 µg/mL] | 27.89 [10 µg/mL] | -1.36 | -4.08 | -16.31 | -5.41 | Inf | 1.90 |
| CBD | CBD-A | 3.18 [1 µg/mL] | 41.84 [15 µg/mL] | -1.53 | -3.23 | -13.56 | -4.52 | Inf | 5.10 |
| CBD | CBD-A | 3.18 [1 µg/mL] | 55.79 [20 µg/mL] | -1.61 | -1.43 | -16.82 | -2.71 | 1.43 | 7.83 |
| CBD | CBD-A | 3.18 [1 µg/mL] | 83.68 [30 µg/mL] | 3.79 | 8.09 | -9.80 | 7.04 | 1.11 | 36.78 |
| CBD | CBD-A | 3.18 [1 µg/mL] | 111.58 [40 µg/mL] | -3.58 | -6.40 | -36.43 | -6.99 | 1.24 | 51.05 |
| CBD | CBD-A | 6.36 [2 µg/mL] | 0.00 [0 µg/mL] | 0.00 | 0.00 | 0.00 | 0.00 | / | 3.10 |
| CBD | CBD-A | 6.36 [2 µg/mL] | 13.95 [5 µg/mL] | -2.14 | -7.43 | -21.81 | -8.90 | Inf | -4.44 |
| CBD | CBD-A | 6.36 [2 µg/mL] | 27.89 [10 µg/mL] | -0.59 | -5.84 | -18.59 | -8.62 | Inf | 0.22 |
| CBD | CBD-A | 6.36 [2 µg/mL] | 41.84 [15 µg/mL] | 0.88 | 2.02 | -17.91 | -0.76 | 1.41 | 9.79 |
| CBD | CBD-A | 6.36 [2 µg/mL] | 55.79 [20 µg/mL] | 5.05 | 7.97 | -16.76 | 5.22 | 1.29 | 17.84 |
| CBD | CBD-A | 6.36 [2 µg/mL] | 83.68 [30 µg/mL] | 18.74 | 19.25 | -25.43 | 17.08 | 1.18 | 47.93 |
| CBD | CBD-A | 6.36 [2 µg/mL] | 111.58 [40 µg/mL] | 19.88 | 20.74 | -41.07 | 19.47 | 1.09 | 76.45 |
| CBD | CBD-A | 9.54 [3 µg/mL] | 0.00 [0 µg/mL] | 0.00 | 0.00 | 0.00 | 0.00 | / | 5.93 |
| CBD | CBD-A | 9.54 [3 µg/mL] | 13.95 [5 µg/mL] | -3.52 | -6.43 | -19.83 | -8.72 | Inf | -1.07 |
| CBD | CBD-A | 9.54 [3 µg/mL] | 27.89 [10 µg/mL] | -2.12 | -0.87 | -23.34 | -5.49 | Inf | 5.69 |
| CBD | CBD-A | 9.54 [3 µg/mL] | 41.84 [15 µg/mL] | 1.90 | 5.11 | -31.48 | -0.14 | 1.43 | 14.95 |
| CBD | CBD-A | 9.54 [3 µg/mL] | 55.79 [20 µg/mL] | 17.87 | 23.98 | -28.27 | 18.73 | 1.24 | 34.75 |
| CBD | CBD-A | 9.54 [3 µg/mL] | 83.68 [30 µg/mL] | 55.03 | 60.01 | -14.91 | 55.88 | 1.00 | 87.34 |

| | | | | | | | | | |
|---|---|---|---|---|---|---|---|---|---|
| CBD | CBD-A | 9.54 [3 µg/mL] | 111.58 [40 µg/mL] | 40.50 | 42.22 | -66.39 | 39.78 | 1.07 | 100.04 |
| CBD | CBD-A | 12.72 [4 µg/mL] | 0.00 [0 µg/mL] | 0.00 | 0.00 | 0.00 | 0.00 | / | 13.22 |
| CBD | CBD-A | 12.72 [4 µg/mL] | 13.95 [5 µg/mL] | -3.34 | -0.40 | -17.06 | -2.52 | 1.26 | 12.47 |
| CBD | CBD-A | 12.72 [4 µg/mL] | 27.89 [10 µg/mL] | 0.64 | 6.16 | -27.09 | 0.84 | 1.31 | 19.74 |
| CBD | CBD-A | 12.72 [4 µg/mL] | 41.84 [15 µg/mL] | 11.16 | 17.61 | -42.68 | 10.29 | 1.31 | 30.94 |
| CBD | CBD-A | 12.72 [4 µg/mL] | 55.79 [20 µg/mL] | 32.10 | 39.11 | -44.43 | 30.88 | 1.24 | 52.18 |
| CBD | CBD-A | 12.72 [4 µg/mL] | 83.68 [30 µg/mL] | 56.43 | 68.14 | -56.41 | 58.82 | 1.07 | 96.80 |
| CBD | CBD-A | 12.72 [4 µg/mL] | 111.58 [40 µg/mL] | 39.05 | 44.13 | -121.17 | 38.65 | 1.21 | 101.68 |
| CBD | CBD-A | 15.9 [5 µg/mL] | 0.00 [0 µg/mL] | 0.00 | 0.00 | 0.00 | 0.00 | / | 34.84 |
| CBD | CBD-A | 15.9 [5 µg/mL] | 13.95 [5 µg/mL] | -0.26 | 0.64 | -25.55 | -1.01 | 1.15 | 37.68 |
| CBD | CBD-A | 15.9 [5 µg/mL] | 27.89 [10 µg/mL] | 11.73 | 16.70 | -32.20 | 12.68 | 1.17 | 50.96 |
| CBD | CBD-A | 15.9 [5 µg/mL] | 41.84 [15 µg/mL] | 21.92 | 22.79 | -55.75 | 17.22 | 1.25 | 57.51 |
| CBD | CBD-A | 15.9 [5 µg/mL] | 55.79 [20 µg/mL] | 40.25 | 48.54 | -57.30 | 42.28 | 1.16 | 81.45 |
| CBD | CBD-A | 15.9 [5 µg/mL] | 83.68 [30 µg/mL] | 45.48 | 65.21 | -111.57 | 47.05 | 1.21 | 100.24 |
| CBD | CBD-A | 15.9 [5 µg/mL] | 111.58 [40 µg/mL] | 29.95 | 42.07 | -185.07 | 27.92 | 1.37 | 99.82 |

**Table S3: Drug concentrations (µg/mL and µM). synergy scores and CI for the CBD-BCP combination in MDA-MB-231 cells**

Inf: infinity - /: not calculated – CI: Combination Index (CI < 1, CI=1, and CI > 1 indicate synergy, additivity and antagonism, respectively). Response % corresponds to the % of inhibition of MDA-MB-231 cell vitality after 72 hours of treatment (represented by the mean of three independent experiment).

| Drug 1 | Drug 2 | Drug 1 µM - [µg/mL] | Drug 2 µM - [µg/mL] | ZIP synergy score | HSA synergy score | Loewe synergy score | Bliss synergy score | CI | Response % |
|---|---|---|---|---|---|---|---|---|---|
| CBD | BCP | 0.00 [0 µg/mL] | 0.00 [0 µg/mL] | 0.00 | 0.00 | 0.00 | 0.00 | / | 0.00 |
| CBD | BCP | 0.00 [0 µg/mL] | 24.47 [5 µg/mL] | 0.00 | 0.00 | 0.00 | 0.00 | / | 4.07 |
| CBD | BCP | 0.00 [0 µg/mL] | 48.94 [10 µg/mL] | 0.00 | 0.00 | 0.00 | 0.00 | / | 8.26 |
| CBD | BCP | 0.00 [0 µg/mL] | 97.87 [20 µg/mL] | 0.00 | 0.00 | 0.00 | 0.00 | / | 13.56 |
| CBD | BCP | 0.00 [0 µg/mL] | 146.81 [30 µg/mL] | 0.00 | 0.00 | 0.00 | 0.00 | / | 44.81 |
| CBD | BCP | 0.00 [0 µg/mL] | 195.74 [40 µg/mL] | 0.00 | 0.00 | 0.00 | 0.00 | / | 63.39 |
| CBD | BCP | 3.18 [1 µg/mL] | 0.00 [0 µg/mL] | 0.00 | 0.00 | 0.00 | 0.00 | / | 1.74 |
| CBD | BCP | 3.18 [1 µg/mL] | 24.47 [5 µg/mL] | -2.28 | -3.76 | -15.36 | -5.56 | Inf | 0.38 |
| CBD | BCP | 3.18 [1 µg/mL] | 48.94 [10 µg/mL] | -0.23 | -6.35 | -13.10 | -8.08 | Inf | 2.83 |
| CBD | BCP | 3.18 [1 µg/mL] | 97.87 [20 µg/mL] | -3.49 | -3.42 | -21.53 | -5.05 | 1.42 | 10.35 |
| CBD | BCP | 3.18 [1 µg/mL] | 146.81 [30 µg/mL] | 31.46 | 32.28 | 19.33 | 31.21 | 0.37 | 76.67 |
| CBD | BCP | 3.18 [1 µg/mL] | 195.74 [40 µg/mL] | 33.21 | 35.84 | 30.29 | 35.14 | 0.39 | 98.75 |
| CBD | BCP | 6.36 [2 µg/mL] | 0.00 [0 µg/mL] | 0.00 | 0.00 | 0.00 | 0.00 | / | 3.10 |
| CBD | BCP | 6.36 [2 µg/mL] | 24.47 [5 µg/mL] | -2.02 | -3.93 | -15.46 | -7.00 | Inf | 0.07 |
| CBD | BCP | 6.36 [2 µg/mL] | 48.94 [10 µg/mL] | 7.68 | 3.66 | -3.09 | 0.66 | 1.06 | 12.80 |
| CBD | BCP | 6.36 [2 µg/mL] | 97.87 [20 µg/mL] | 56.92 | 60.63 | 27.99 | 57.80 | 0.46 | 67.23 |
| CBD | BCP | 6.36 [2 µg/mL] | 146.81 [30 µg/mL] | 43.20 | 41.89 | 21.88 | 40.07 | 0.50 | 86.54 |
| CBD | BCP | 6.36 [2 µg/mL] | 195.74 [40 µg/mL] | 29.75 | 27.61 | 12.54 | 26.40 | 0.55 | 89.50 |
| CBD | BCP | 7.95 [2.5 µg/mL] | 0.00 [0 µg/mL] | 0.00 | 0.00 | 0.00 | 0.00 | / | 3.52 |
| CBD | BCP | 7.95 [2.5 µg/mL] | 24.47 [5 µg/mL] | -2.13 | 2.65 | -8.96 | -0.70 | 1.13 | 6.50 |
| CBD | BCP | 7.95 [2.5 µg/mL] | 48.94 [10 µg/mL] | 18.85 | 20.00 | 6.15 | 16.82 | 0.91 | 28.16 |
| CBD | BCP | 7.95 [2.5 µg/mL] | 97.87 [20 µg/mL] | 70.92 | 73.68 | 31.60 | 70.67 | 0.51 | 89.03 |
| CBD | BCP | 7.95 [2.5 µg/mL] | 146.81 [30 µg/mL] | 47.77 | 50.03 | 24.03 | 48.12 | 0.55 | 93.38 |
| CBD | BCP | 7.95 [2.5 µg/mL] | 195.74 [40 µg/mL] | 33.23 | 33.05 | 12.24 | 31.76 | 0.61 | 95.71 |
| CBD | BCP | 9.54 [3 µg/mL] | 0.00 [0 µg/mL] | 0.00 | 0.00 | 0.00 | 0.00 | / | 5.93 |
| CBD | BCP | 9.54 [3 µg/mL] | 24.47 [5 µg/mL] | -3.31 | -3.06 | -12.94 | -6.84 | Inf | 3.12 |
| CBD | BCP | 9.54 [3 µg/mL] | 48.94 [10 µg/mL] | 25.82 | 27.51 | 3.28 | 22.28 | 0.94 | 34.67 |

| | | | | | | | | | |
|---|---|---|---|---|---|---|---|---|---|
| CBD | BCP | 9.54 [3 µg/mL] | 97.87 [20 µg/mL] | 75.59 | 81.88 | 35.25 | 76.93 | 0.57 | 95.34 |
| CBD | BCP | 9.54 [3 µg/mL] | 146.81 [30 µg/mL] | 49.18 | 51.24 | 21.71 | 48.10 | 0.63 | 96.74 |
| CBD | BCP | 9.54 [3 µg/mL] | 195.74 [40 µg/mL] | 33.65 | 36.99 | 0.57 | 34.87 | 0.68 | 99.56 |
| CBD | BCP | 12.72 [4 µg/mL] | 0.00 [0 µg/mL] | 0.00 | 0.00 | 0.00 | 0.00 | / | 13.22 |
| CBD | BCP | 12.72 [4 µg/mL] | 24.47 [5 µg/mL] | 19.99 | 25.07 | 4.44 | 21.57 | 0.94 | 37.85 |
| CBD | BCP | 12.72 [4 µg/mL] | 48.94 [10 µg/mL] | 44.23 | 45.36 | 4.30 | 37.65 | 0.94 | 58.10 |
| CBD | BCP | 12.72 [4 µg/mL] | 97.87 [20 µg/mL] | 65.89 | 78.98 | 20.65 | 68.36 | 0.72 | 91.65 |
| CBD | BCP | 12.72 [4 µg/mL] | 146.81 [30 µg/mL] | 46.80 | 53.56 | -1.10 | 46.59 | 0.77 | 98.50 |
| CBD | BCP | 12.72 [4 µg/mL] | 195.74 [40 µg/mL] | 34.25 | 40.12 | -15.28 | 35.45 | 0.82 | 103.71 |
| CBD | BCP | 15.9 [5 µg/mL] | 0.00 [0 µg/mL] | 0.00 | 0.00 | 0.00 | 0.00 | / | 34.84 |
| CBD | BCP | 15.9 [5 µg/mL] | 24.47 [5 µg/mL] | 53.53 | 56.28 | 32.16 | 53.57 | 0.79 | 88.99 |
| CBD | BCP | 15.9 [5 µg/mL] | 48.94 [10 µg/mL] | 56.62 | 59.47 | 19.16 | 53.44 | 0.81 | 92.08 |
| CBD | BCP | 15.9 [5 µg/mL] | 97.87 [20 µg/mL] | 54.44 | 65.12 | -1.51 | 55.70 | 0.86 | 96.89 |
| CBD | BCP | 15.9 [5 µg/mL] | 146.81 [30 µg/mL] | 37.88 | 54.94 | -18.03 | 37.75 | 0.91 | 100.70 |
| CBD | BCP | 15.9 [5 µg/mL] | 195.74 [40 µg/mL] | 26.61 | 39.28 | -35.17 | 27.40 | 0.97 | 102.11 |

**Table S4: Drug concentrations (µg/mL and µM). synergy scores and CI for the three-drug combination (CBD+CBG+BCP) in MDA-MB-231 cells**

Inf: infinity - /: not calculated – CI: Combination Index (CI < 1, CI=1, and CI > 1 indicate synergy, additivity and antagonism, respectively). Response % corresponds to the % of inhibition of MDA-MB-231 cell vitality after 72 hours of treatment (represented by the mean of three independent experiment).

| Drug 1 | Drug 2 | Drug 3 | Drug 1 µM - [µg/mL] | Drug 2 µM - [µg/mL] | Drug 3 µM - [µg/mL] | ZIP synergy score | HSA synergy score | Loewe synergy score | Bliss synergy score | CI | Response % |
|---|---|---|---|---|---|---|---|---|---|---|---|
| CBD | CBG | BCP | 0 [0 µg/mL] | 0 [0 µg/mL] | 0 [0 µg/mL] | 0.00 | 0.00 | 0.00 | 0.00 | / | 0.00 |
| CBD | CBG | BCP | 3.18 [1 µg/mL] | 0 [0 µg/mL] | 0 [0 µg/mL] | 0.00 | 0.00 | 0.00 | 0.00 | / | 1.74 |
| CBD | CBG | BCP | 9.54 [3 µg/mL] | 0 [0 µg/mL] | 0 [0 µg/mL] | 0.00 | 0.00 | 0.00 | 0.00 | / | 5.93 |
| CBD | CBG | BCP | 15.9 [5 µg/mL] | 0 [0 µg/mL] | 0 [0 µg/mL] | 0.00 | 0.00 | 0.00 | 0.00 | / | 34.84 |
| CBD | CBG | BCP | 0 [0 µg/mL] | 9.48 [3 µg/mL] | 0 [0 µg/mL] | 0.00 | 0.00 | 0.00 | 0.00 | / | -2.52 |
| CBD | CBG | BCP | 3.18 [1 µg/mL] | 9.48 [3 µg/mL] | 0 [0 µg/mL] | 2.02 | 0.96 | -1.52 | 3.53 | 1.02 | 2.35 |
| CBD | CBG | BCP | 9.54 [3 µg/mL] | 9.48 [3 µg/mL] | 0 [0 µg/mL] | 19.58 | 18.37 | -12.62 | 20.82 | 1.11 | 22.89 |
| CBD | CBG | BCP | 15.9 [5 µg/mL] | 9.48 [3 µg/mL] | 0 [0 µg/mL] | 50.41 | 49.74 | 31.17 | 51.44 | 0.75 | 85.86 |
| CBD | CBG | BCP | 0 [0 µg/mL] | 18.96 [6 µg/mL] | 0 [0 µg/mL] | 0.00 | 0.00 | 0.00 | 0.00 | / | 5.91 |
| CBD | CBG | BCP | 3.18 [1 µg/mL] | 18.96 [6 µg/mL] | 0 [0 µg/mL] | 15.06 | 16.03 | -18.14 | 14.52 | 1.13 | 21.54 |
| CBD | CBG | BCP | 9.54 [3 µg/mL] | 18.96 [6 µg/mL] | 0 [0 µg/mL] | 68.30 | 73.38 | 31.37 | 68.64 | 0.46 | 79.52 |
| CBD | CBG | BCP | 15.9 [5 µg/mL] | 18.96 [6 µg/mL] | 0 [0 µg/mL] | 64.88 | 68.29 | 49.33 | 64.92 | 0.69 | 102.31 |
| CBD | CBG | BCP | 0 [0 µg/mL] | 28.44 [9 µg/mL] | 0 [0 µg/mL] | 0.00 | 0.00 | 0.00 | 0.00 | / | 48.97 |
| CBD | CBG | BCP | 3.18 [1 µg/mL] | 28.44 [9 µg/mL] | 0 [0 µg/mL] | 19.22 | 19.74 | 21.57 | 18.94 | 0.16 | 68.43 |
| CBD | CBG | BCP | 9.54 [3 µg/mL] | 28.44 [9 µg/mL] | 0 [0 µg/mL] | 44.88 | 47.88 | 44.93 | 44.78 | 0.42 | 96.14 |
| CBD | CBG | BCP | 15.9 [5 µg/mL] | 28.44 [9 µg/mL] | 0 [0 µg/mL] | 36.03 | 53.27 | 49.74 | 35.91 | 0.69 | 103.54 |
| CBD | CBG | BCP | 0 [0 µg/mL] | 0 [0 µg/mL] | 24.47 [5 µg/mL] | 0.00 | 0.00 | 0.00 | 0.00 | / | 4.07 |
| CBD | CBG | BCP | 3.18 [1 µg/mL] | 0 [0 µg/mL] | 24.47 [5 µg/mL] | -0.98 | -3.54 | -5.75 | -5.06 | / | 0.47 |
| CBD | CBG | BCP | 9.54 [3 µg/mL] | 0 [0 µg/mL] | 24.47 [5 µg/mL] | -2.79 | -1.05 | -9.26 | -4.88 | 1.80 | 4.46 |
| CBD | CBG | BCP | 15.9 [5 µg/mL] | 0 [0 µg/mL] | 24.47 [5 µg/mL] | 26.85 | 28.51 | 21.47 | 25.83 | 0.86 | 70.55 |
| CBD | CBG | BCP | 0 [0 µg/mL] | 9.48 [3 µg/mL] | 24.47 [5 µg/mL] | -6.19 | -15.28 | -19.27 | -12.77 | / | -10.89 |
| CBD | CBG | BCP | 3.18 [1 µg/mL] | 9.48 [3 µg/mL] | 24.47 [5 µg/mL] | -5.61 | -15.14 | -23.87 | -14.19 | Inf | -9.76 |

| | | | | | | | | | | | |
|---|---|---|---|---|---|---|---|---|---|---|---|
| CBD | CBG | BCP | 9.54 [3 μg/mL] | 9.48 [3 μg/mL] | 24.47 [5 μg/mL] | 3.26 | -1.99 | -37.73 | -3.47 | Inf | 4.25 |
| CBD | CBG | BCP | 15.9 [5 μg/mL] | 9.48 [3 μg/mL] | 24.47 [5 μg/mL] | 26.96 | 13.86 | -4.71 | 12.82 | 1.21 | 47.49 |
| CBD | CBG | BCP | 0 [0 μg/mL] | 18.96 [6 μg/mL] | 24.47 [5 μg/mL] | -7.78 | -10.48 | -27.67 | -14.19 | / | -2.65 |
| CBD | CBG | BCP | 3.18 [1 μg/mL] | 18.96 [6 μg/mL] | 24.47 [5 μg/mL] | 10.50 | 13.41 | -26.07 | 8.26 | 1.29 | 18.69 |
| CBD | CBG | BCP | 9.54 [3 μg/mL] | 18.96 [6 μg/mL] | 24.47 [5 μg/mL] | 57.34 | 61.68 | 19.22 | 53.31 | 0.51 | 67.02 |
| CBD | CBG | BCP | 15.9 [5 μg/mL] | 18.96 [6 μg/mL] | 24.47 [5 μg/mL] | 57.93 | 63.37 | 44.41 | 57.46 | 0.72 | 93.71 |
| CBD | CBG | BCP | 0 [0 μg/mL] | 28.44 [9 μg/mL] | 24.47 [5 μg/mL] | 0.44 | 2.93 | 4.76 | 0.89 | 0.59 | 54.76 |
| CBD | CBG | BCP | 3.18 [1 μg/mL] | 28.44 [9 μg/mL] | 24.47 [5 μg/mL] | 17.37 | 18.87 | 20.71 | 16.07 | 0.19 | 69.28 |
| CBD | CBG | BCP | 9.54 [3 μg/mL] | 28.44 [9 μg/mL] | 24.47 [5 μg/mL] | 40.69 | 44.56 | 41.61 | 39.55 | 0.45 | 96.36 |
| CBD | CBG | BCP | 15.9 [5 μg/mL] | 28.44 [9 μg/mL] | 24.47 [5 μg/mL] | 31.79 | 48.39 | 44.86 | 29.70 | 0.73 | 97.23 |
| CBD | CBG | BCP | 3.18 [1 μg/mL] | 0 [0 μg/mL] | 48.94 [10 μg/mL] | -3.02 | -4.44 | -8.52 | -5.92 | 4.11 | 2.85 |
| CBD | CBG | BCP | 9.54 [3 μg/mL] | 0 [0 μg/mL] | 48.94 [10 μg/mL] | 12.89 | 17.49 | 4.95 | 12.22 | 0.95 | 29.26 |
| CBD | CBG | BCP | 15.9 [5 μg/mL] | 0 [0 μg/mL] | 48.94 [10 μg/mL] | 46.64 | 50.57 | 38.59 | 45.99 | 0.79 | 85.60 |
| CBD | CBG | BCP | 0 [0 μg/mL] | 9.48 [3 μg/mL] | 48.94 [10 μg/mL] | -8.07 | -15.37 | -21.19 | -12.94 | / | -7.32 |
| CBD | CBG | BCP | 3.18 [1 μg/mL] | 9.48 [3 μg/mL] | 48.94 [10 μg/mL] | -5.05 | -5.63 | -17.89 | -4.71 | Inf | 0.08 |
| CBD | CBG | BCP | 9.54 [3 μg/mL] | 9.48 [3 μg/mL] | 48.94 [10 μg/mL] | 13.59 | 12.75 | -22.75 | 9.77 | 2.17 | 22.20 |
| CBD | CBG | BCP | 15.9 [5 μg/mL] | 9.48 [3 μg/mL] | 48.94 [10 μg/mL] | 32.88 | 18.30 | -3.36 | 15.30 | 1.13 | 52.67 |
| CBD | CBG | BCP | 0 [0 μg/mL] | 18.96 [6 μg/mL] | 48.94 [10 μg/mL] | -6.97 | -3.12 | -25.84 | -7.63 | / | 6.30 |
| CBD | CBG | BCP | 3.18 [1 μg/mL] | 18.96 [6 μg/mL] | 48.94 [10 μg/mL] | 19.67 | 25.15 | -12.70 | 19.24 | 1.18 | 30.45 |
| CBD | CBG | BCP | 9.54 [3 μg/mL] | 18.96 [6 μg/mL] | 48.94 [10 μg/mL] | 67.70 | 77.38 | 34.23 | 67.71 | 0.50 | 84.37 |
| CBD | CBG | BCP | 15.9 [5 μg/mL] | 18.96 [6 μg/mL] | 48.94 [10 μg/mL] | 57.43 | 65.54 | 43.49 | 57.83 | 0.74 | 98.90 |
| CBD | CBG | BCP | 0 [0 μg/mL] | 28.44 [9 μg/mL] | 48.94 [10 μg/mL] | 2.99 | 6.39 | 8.22 | 2.90 | 0.30 | 56.54 |
| CBD | CBG | BCP | 3.18 [1 μg/mL] | 28.44 [9 μg/mL] | 48.94 [10 μg/mL] | 24.99 | 28.81 | 30.65 | 24.57 | 0.20 | 77.79 |
| CBD | CBG | BCP | 9.54 [3 μg/mL] | 28.44 [9 μg/mL] | 48.94 [10 μg/mL] | 43.30 | 49.97 | 47.02 | 43.59 | 0.46 | 99.57 |
| CBD | CBG | BCP | 15.9 [5 μg/mL] | 28.44 [9 μg/mL] | 48.94 [10 μg/mL] | 32.18 | 52.31 | 45.68 | 32.66 | 0.73 | 101.99 |
| CBD | CBG | BCP | 3.18 [1 μg/mL] | 0 [0 μg/mL] | 97.87 [20 μg/mL] | -3.44 | -5.90 | -9.79 | -7.27 | 3.24 | 8.63 |
| CBD | CBG | BCP | 9.54 [3 μg/mL] | 0 [0 μg/mL] | 97.87 [20 μg/mL] | 62.60 | 66.89 | 50.61 | 61.52 | 0.55 | 75.37 |
| CBD | CBG | BCP | 15.9 [5 μg/mL] | 0 [0 μg/mL] | 97.87 [20 μg/mL] | 53.40 | 61.50 | 39.30 | 52.57 | 0.79 | 95.38 |
| CBD | CBG | BCP | 0 [0 μg/mL] | 9.48 [3 μg/mL] | 97.87 [20 μg/mL] | -6.74 | -9.93 | -18.72 | -7.68 | 4.96 | 3.01 |
| CBD | CBG | BCP | 3.18 [1 μg/mL] | 9.48 [3 μg/mL] | 97.87 [20 μg/mL] | 18.90 | 18.62 | 2.34 | 19.47 | 0.92 | 32.05 |

| | | | | | | | | | | | |
|---|---|---|---|---|---|---|---|---|---|---|---|
| CBD | CBG | BCP | 9.54 [3 µg/mL] | 9.48 [3 µg/mL] | 97.87 [20 µg/mL] | 59.00 | 59.27 | 28.12 | 56.02 | 0.59 | 71.97 |
| CBD | CBG | BCP | 15.9 [5 µg/mL] | 9.48 [3 µg/mL] | 97.87 [20 µg/mL] | 54.58 | 65.47 | 34.91 | 58.01 | 0.78 | 99.73 |
| CBD | CBG | BCP | 0 [0 µg/mL] | 18.96 [6 µg/mL] | 97.87 [20 µg/mL] | 34.46 | 39.40 | 16.72 | 34.96 | 0.48 | 53.72 |
| CBD | CBG | BCP | 3.18 [1 µg/mL] | 18.96 [6 µg/mL] | 97.87 [20 µg/mL] | 61.21 | 66.26 | 34.84 | 60.52 | 0.25 | 84.86 |
| CBD | CBG | BCP | 9.54 [3 µg/mL] | 18.96 [6 µg/mL] | 97.87 [20 µg/mL] | 75.82 | 85.70 | 47.94 | 76.17 | 0.50 | 99.39 |
| CBD | CBG | BCP | 15.9 [5 µg/mL] | 18.96 [6 µg/mL] | 97.87 [20 µg/mL] | 54.58 | 66.86 | 35.90 | 55.02 | 0.77 | 101.54 |
| CBD | CBG | BCP | 0 [0 µg/mL] | 28.44 [9 µg/mL] | 97.87 [20 µg/mL] | 39.23 | 46.42 | 48.25 | 39.58 | 0.08 | 96.34 |
| CBD | CBG | BCP | 3.18 [1 µg/mL] | 28.44 [9 µg/mL] | 97.87 [20 µg/mL] | 42.04 | 49.21 | 49.20 | 41.69 | 0.22 | 99.01 |
| CBD | CBG | BCP | 9.54 [3 µg/mL] | 28.44 [9 µg/mL] | 97.87 [20 µg/mL] | 41.99 | 51.90 | 48.96 | 42.39 | 0.50 | 102.06 |
| CBD | CBG | BCP | 15.9 [5 µg/mL] | 28.44 [9 µg/mL] | 97.87 [20 µg/mL] | 30.08 | 52.29 | 36.77 | 30.49 | 0.77 | 102.63 |
