## Supplementary Figures for "Synergistic induction of a lethal Autosis-to-Apoptosis switch by Phytocannabinoids and beta-Caryophyllene in Triple-Negative Breast Cancer Cells"

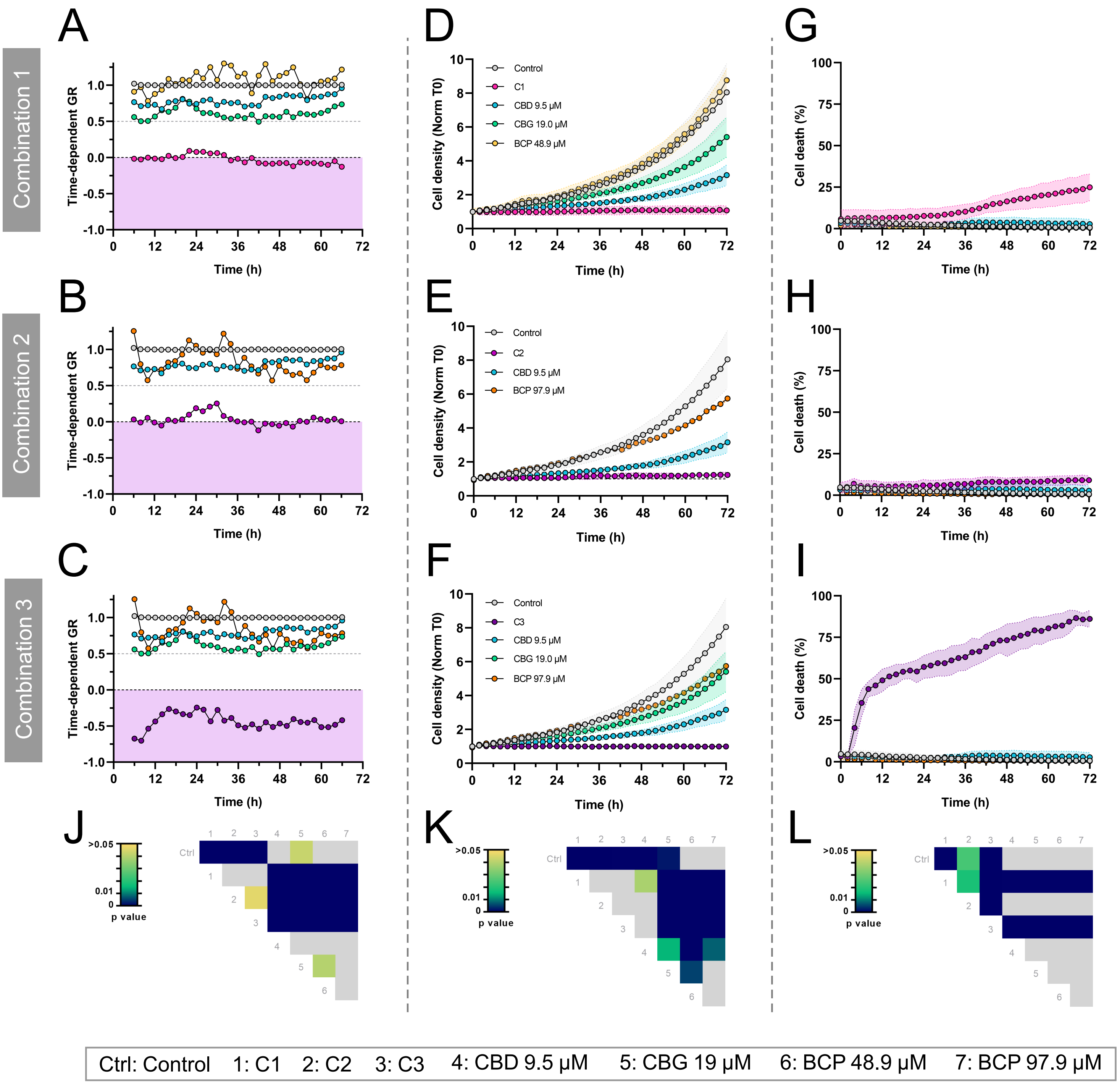


**Supplementary Figure S1. Kinetic analysis of the top synergistic combinations on the growth of Hs578T cells.** Cell responses were analyzed by live-cell imaging for 72h, and automated image analysis was performed to quantify the time-dependent GR values **(A-C)**, the evolution of cell density **(D-F)** and cell death **(G-I)**. Data are expressed as mean + SD from three independent experiments (in triplicate). Statistical analyses were performed using one-way ANOVA followed by Tukey's post hoc test for multiple comparisons. A heatmap summarizing the multiple comparisons is provided below each graph. **(J)** statistical analysis on the GR value after 72h of treatment. **(K-L)** Statistical analysis was performed on the area under the curve (AUC) of each individual curve. Statistical significance was set at $p < 0.05$. Grey colour corresponds to non-significant p-values.

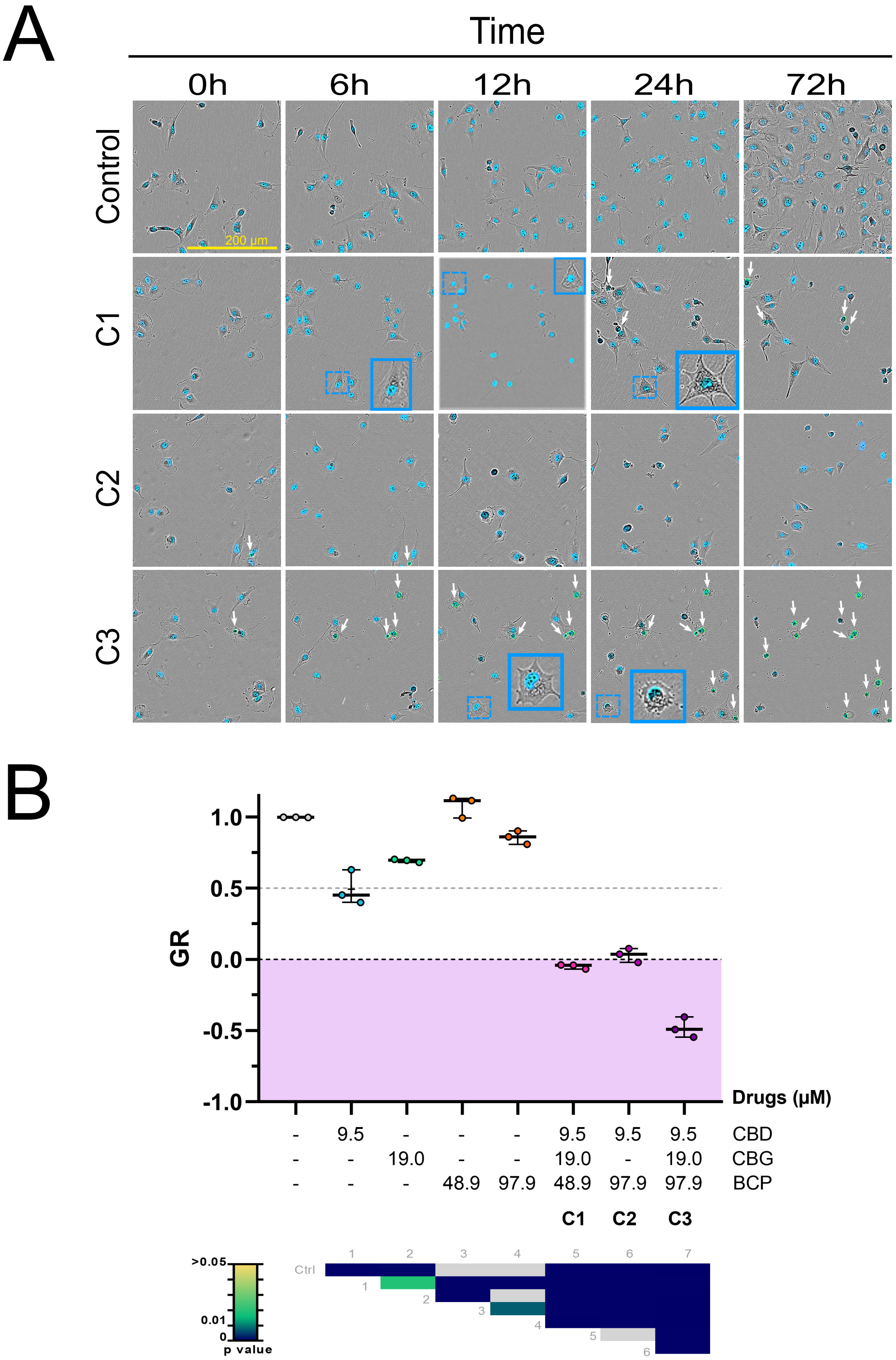


**Supplementary Figure S2. Live-cell imaging reveals morphodynamic alterations induced by synergistic combinations in Hs578T cells.** Hs578T cells were treated with vehicle (0.3% DMSO - Control) or combinations C1 to C3 as described in Table 1 in the presence of SYTOX Green and SPY-DNA. **(A)** Automated live-cell imaging was performed for 72h and representative composite images combining the phase contrast, green (SYTOX Green, dead cells), and NiR (SPY650-DNA, nuclei) fluorescence channels obtained at different time points are displayed. Dead cells (SPY+/SYTOX Green+ nuclei) are indicated by white arrows. Insets show higher magnifications of specific elements. Blue boxes highlight cells exhibiting cytoplasmic vacuoles. Scale bar corresponds to 200 µm. **(B)** Growth rate (GR) values were calculated for each combination on Hs578T. Data represents the median of four independent experiments and was analyzed using one-way ANOVA followed by Tukey's post hoc test for multiple comparisons. A heatmap summarizing the multiple comparisons is provided below the graph. Ctrl: control; 1 to 7 corresponds to the experimental conditions in the same order as listed in the figure. Statistical significance was set at $p < 0.05$. Grey colour corresponds to non-significant p-values.

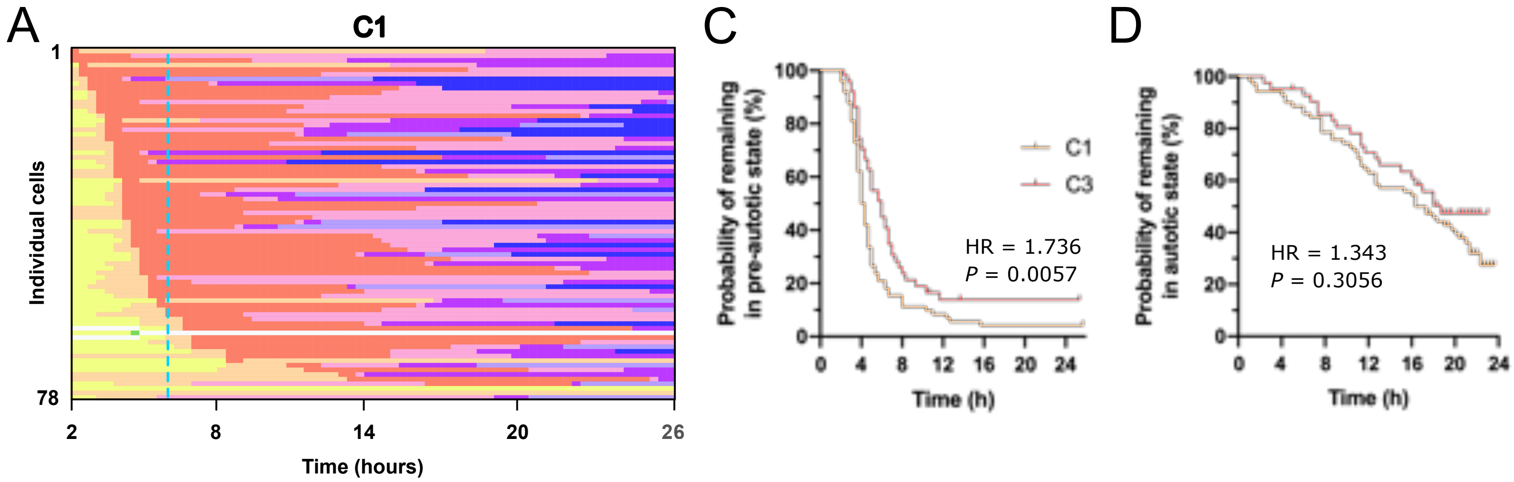


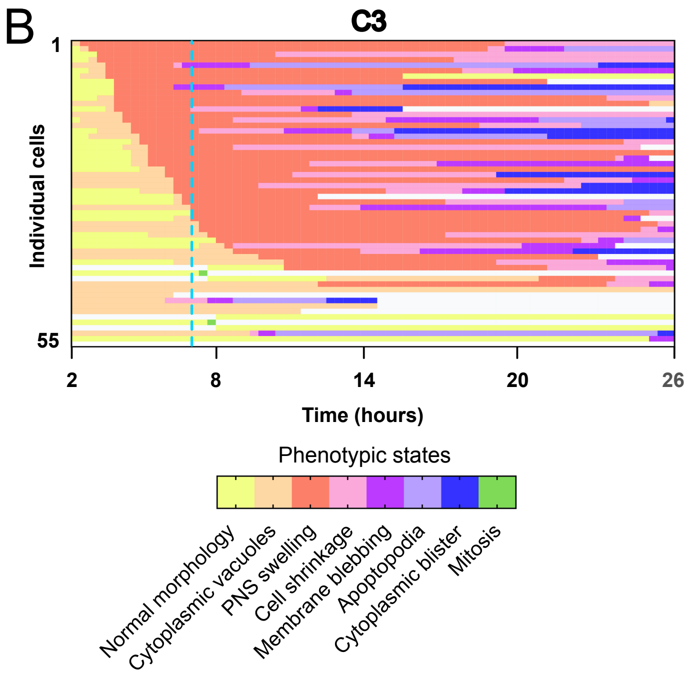


E

| | C1 | C3 |
|---|---|---|
| Autosis penetrance | 92.3% | 82.4% |
| Median autosis entry time (IQR) | 6h (5.3–7.0) | 7h (5.7–8.7) |
| Transition from autosis to apoptotic blebbing | 68.1% | 50.0% |
| Median lag time between PNS swelling and membrane blebbing (IQR) | 11.3h (7.7–16.8) | 11.3h (7.3–16.0) |
| Direct transition from autosis to necrosis | 5.6% | 4.8% |
| Mitosis | 1.3% | 3.9% |

**Supplementary Figure S3. Comparative single-cell kinetic mapping of C1- and C3-induced cell death.** MDA-MB-231 cells were monitored via label-free holotomographic microscopy at 20-minute resolution for 24 h. Imaging commenced 2 h post-treatment to resolve the subcellular phenotypic trajectories. **(A–B)** Single-cell phenotypic heatmaps for C1-treated (n=78) and C3-treated (n=55) lineages. Each horizontal track represents the temporal evolution of a single cell, with colours denoting discrete phenotypic states. Trajectories are synchronized by the onset of perinuclear space (PNS) swelling. Vertical blue dotted lines indicate the median initiation time for autotic commitment (6.0 h for C1; 7.0 h for C3). **(C)** Kinetics of autosis initiation. Kaplan-Meier plot illustrating the time-to-onset of the autotic program, defined by the first appearance of focal PNS swelling. The Y-axis represents the probability of remaining in a pre-autotic state (cells yet to exhibit PNS swelling) over time. C1-treated cells exhibited a significantly faster initiation compared to the C3 (Log-rank Mantel-Cox test, $P < 0.05$). **(D)** Temporal conservation of the execution phase. Kaplan-Meier plot showing the execution lag, defined as the interval between the onset of autotic commitment (PNS swelling) and the initiation of terminal apoptotic blebbing. The Y-axis represents the probability of autotic survival (the proportion of committed cells that have not yet transitioned to the execution phase). Both treatments showed similar conversion efficiencies (Log-rank Mantel-Cox test, $P = 0.3056$). **(E)** Comparison of the major phenotypic alterations induced by C1 and C3 treatments.

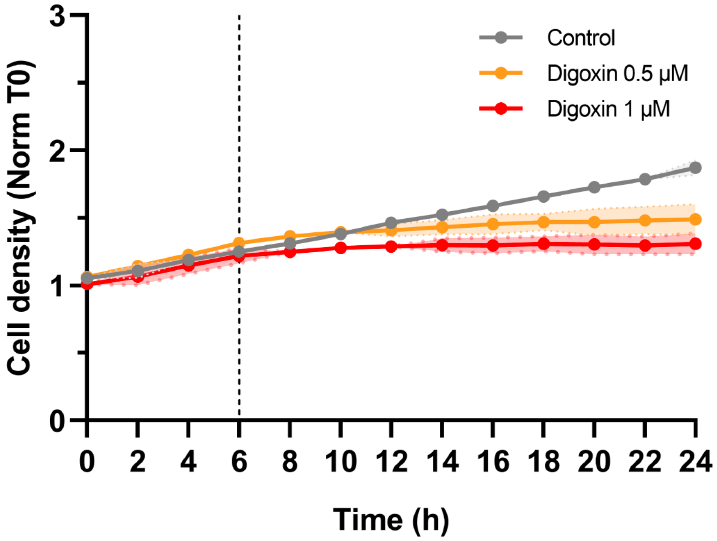


**Supplementary Figure S4. Kinetic monitoring of MDA-MB-231 cell growth under digoxin treatment.** Time-course evolution of SPY+ nuclei density was quantified using automated live-cell imaging with images acquired every 2 hours over a 24 h period. Cells were treated with vehicle (control) or digoxin (0.5 µM and 1 µM). Data represent mean ± SD (n=3). This analysis confirms that at the 6 h time point (used for C1-rescue experiments), digoxin does not induce growth inhibition or cytotoxicity compared to control cells ($p > 0.05$).
